# Intermittent ERK oscillations downstream of FGF in mouse embryonic stem cells

**DOI:** 10.1101/2020.12.14.422687

**Authors:** Dhruv Raina, Fiorella Fabris, Luis G. Morelli, Christian Schröter

**Affiliations:** Department of Systemic Cell Biology, Max Planck Institute of Molecular Physiology, Otto-Hahn-Str. 11, 44227 Dortmund, Germany; Instituto de Investigación en Biomedicina de Buenos Aires (IBioBA)—CONICET—Partner Institute of the Max Planck Society, Polo Científico Tecnológico, Godoy Cruz 2390, C1425FQD Buenos Aires, Argentina; Departamento de Física, FCEyN UBA, Ciudad Universitaria, 1428 Buenos Aires, Argentina

**Author notes:** these authors contributed equally (Dhruv Raina, Fiorella Fabris).

## Abstract

Signal transduction networks process extracellular signals to guide cell fate decisions such as to divide, differentiate, or die. These networks can generate characteristic dynamic activities that are shaped by their cell-type specific architecture. The differentiation of pluripotent cells is controlled by FGF/ERK signaling. However, the dynamic activity of the FGF/ERK signaling network in this context remains unexplored. Here we use live cell sensors in wild type and *Fgf4* mutant mouse embryonic stem cells to measure ERK dynamic activity in single cells, in response to defined ligand concentrations. We find that ERK activity oscillates in embryonic stem cells. Single cells can transit between oscillatory and non-oscillatory behavior, leading to heterogeneous dynamic activities in the population. Oscillations become more prevalent with increasing FGF4 dose, while maintaining a robust characteristic timescale. Our results suggest that FGF/ERK signaling operates in the vicinity of a transition point between oscillatory and non-oscillatory dynamics in embryonic stem cells.

## Introduction

Cells rely on signal transduction networks to process signals from their environment, and to guide decisions such as to divide, differentiate, or die (Koseska and Bastiaens, 2017). These networks can produce dynamic activation patterns even at constant stimuli (Antebi et al., 2017; Santos et al., 2007). Dynamic activity patterns are shaped by the cell-type specific architecture of the signal transduction system.

One of the most critical signal transduction systems during early mammalian embryogenesis relays signals from extracellular fibroblast growth factor 4 (FGF4) through the RAS/RAF/MEK/ERK network (Brewer et al., 2016). The differentiation of extraembryonic primitive endoderm cells in the mouse preimplantation embryo depends on FGF/ERK signaling in a dose-dependent manner (Kang et al., 2013; Krawchuk et al., 2013). Embryonic stem cells (ESCs), clonal cell populations that retain the differentiation potential of inner cell mass cells of the preimplantation embryo, are a tractable model system that recapitulates this dose-dependent function of FGF4 (Raina et al., 2020; Schröter et al., 2015). FGF/ERK signaling is also required for maturation of the epiblast lineage in the embryo (Kang et al., 2017; Ohnishi et al., 2014), and controls the corresponding process of transitioning from naïve to primed pluripotency and lineage commitment in ESCs (Kunath et al., 2007; Molotkov et al., 2017). Both in the embryo and ESCs, FGF/ERK signaling is mostly triggered by paracrine FGF4 ligands (Kang et al., 2013; Krawchuk et al., 2013; Kunath et al., 2007). Despite these well-known functions of FGF/ERK signaling during the differentiation of pluripotent cells, little is known about FGF/ERK signaling dynamics in this developmental context.

Revealing intracellular signal transduction dynamics requires live-cell approaches in single cells. Live-cell ERK activity can be monitored with substrate-based sensors that employ FRET or subcellular localization as read-outs (Komatsu et al., 2011; Regot et al., 2014). Analysis of ERK activity in acutely stimulated ESCs expressing a FRET-based sensor revealed a transient peak of activation that decayed over long timescales (Deathridge et al., 2019). However, the short timescale ERK signaling dynamics in the continuous FGF stimulation regimes required to trigger differentiation of ESCs (Hamilton et al., 2019) remains largely unexplored.

Short timescale ERK dynamics upon continuous stimulation of other receptor tyrosine kinases (RTKs) such as the epidermal growth factor (EGF) receptor have been studied in various cell types, revealing a diversity of behaviors. In many, but not all cell types, ERK activity occurs in pulses (Aoki et al., 2013). In several cell types, the frequency of ERK activity pulses depends on EGF concentration or cell density (Albeck et al., 2013; Aoki et al., 2013). This has led to the suggestion of frequency-modulated encoding of information about extracellular signal levels by the RAS/RAF/MEK/ERK network downstream of the EGF receptor (Albeck et al., 2013). In mammary epithelial cells in contrast, pulses of ERK nuclear translocation have a constant frequency across a range of EGF stimulation levels (Shankaran et al., 2009).

Here we use a translocation-based sensor (Regot et al., 2014) to measure short timescale ERK activity dynamics in single ESCs upon continuous FGF stimulation. We find that ERK activity is pulsatile in ESCs, and develop concepts and analysis methods to quantitatively characterize dynamic signatures of pulsing. ERK activity pulses in ESCs are faster than any previously reported ERK dynamics. Pulses occur with high regularity consistent with an oscillatory behavior in a subset of cells. We detect no pulsing in unstimulated *Fgf4* mutant cells, indicating that ERK pulses are driven by FGF4. Controlling extracellular ligand levels in the mutant background, we show that individual ERK pulses have a duration that is independent of ligand levels. However, the extent of the oscillatory behavior increases with FGF4 dose. Finally, we show that ERK pulsing is more prevalent in the early stages of the cell cycle. Our data suggest that the FGF/ERK signal transduction system in ESCs transits between oscillatory and non-oscillatory behavior.

## Results

### ERK activity is dynamic in ESCs

We first explored ERK activation in single ESCs under constant culture conditions that maintain pluripotency. We stained for phosphorylated ERK (pERK) in cells growing in serum + LIF (S+L) and quantified whole-cell pERK levels. We observed pERK staining in cells growing in serum + LIF which was absent in the presence of the MEK-inhibitor PD0325901 (MEKi) (Fig. 1A, B). pERK staining was more heterogeneous in serum + LIF than in the MEKi control. Almost all cells in serum + LIF had pERK staining values above the range covered by MEKi cells (Fig. 1B).

**Fig. 1.**
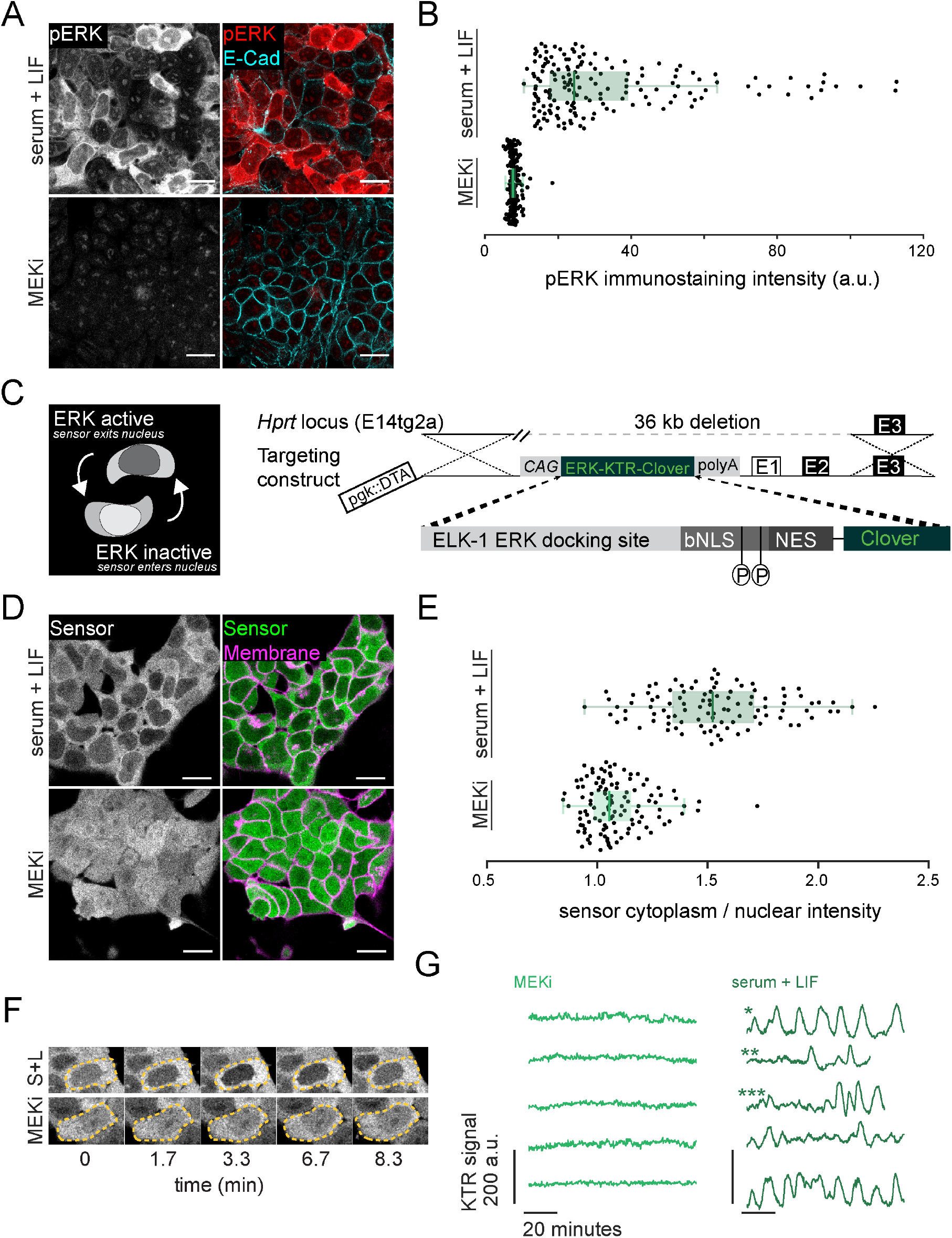
A targeted translocation sensor reveals pulsatile ERK activity in ESCs. **A.** Immunostaining of mESCs growing in serum + LIF medium without (top) or with MEKi (bottom) for pERK and E-Cadherin to mark membranes. The punctate pERK staining within the nucleus is insensitive to MEK inhibition, suggesting it is non-specific. Scale bar = 20 μm. **B.** Quantification of fluorescence staining intensities in single cells stained as in A. n ≥ 100 per condition, green bars indicate medians (Median_S+L_=24.39 a.u., Median_MEKi_=7.75 a.u.; CV_S+L_=0.67, CV_MEKi_=0.17), box bounds are the 25 and 75 percentiles of the distributions, and whiskers are the 5 and 95 percentiles. **C.** Schematic of the ERK-KTR sensor and targeting construct for integration into the Hprt locus. **D.** Subcellular localization of ERK-KTR sensor in live cells in serum + LIF without (top) and with MEKi (bottom). Membranes are stained with live-cell membrane dye CellMaskRed. Scale bar = 20 μm. **E.** Quantification of cytoplasmic to nuclear ratio of sensor fluorescence in single cells imaged as in **D**, green bars indicate medians (Median_S+L_=1.52, Median_MEKi_= 1.05; CV_S+L_=0.18, CV_MEKi_=0.13), box bounds are the 25 and 75 percentiles of the distributions, and whiskers are the 5 and 95 percentiles. **F.** Stills from a movie of ERK-KTR expressing cells growing in serum + LIF without (top) and with MEKi (bottom). Dashed line indicates cell outlines. **G.** Representative traces of the KTR signal obtained as the mean inverted fluorescence intensity within a nuclear ROI in single cells growing in serum + LIF without (right) and with MEKi (left).

The heterogeneous pERK staining in serum + LIF could purely reflect long-term variability between cells as previously reported (Deathridge et al., 2019). In addition, short-term signaling fluctuations could contribute to this variability. To test the extent of short-term signaling fluctuations, we integrated a translocation-based sensor to measure ERK activity in live cells. We generated this cell line by single copy insertion of the ERK-KTR-mClover construct into the *Hprt* open locus (Fig. 1C) to ensure uniformity in expression. Transgenic cells continued to express pluripotency markers (Fig. 1 Supp. 1), and transmitted to the germline of chimeric mice (Simon et al., 2020), indicating that reporter expression does not interfere with pluripotency and differentiation potential.

Phosphorylation of the ERK target site of the sensor leads to its export from the nucleus, thus reporting ERK activity as the cytoplasmic to nuclear (C/N) ratio of reporter localization (Regot et al., 2014) (Fig. 1C). Snapshots of cells growing in serum + LIF showed that the sensor preferentially localized to the cytoplasm, in contrast to the MEKi treated control where it was uniformly distributed (Fig. 1D). Furthermore, the C/N ratio of sensor localization was more variable between cells growing in serum + LIF compared to the MEKi-treated control (Fig. 1E), in line with heterogeneous pERK staining. These qualitative similarities between pERK staining and reporter C/N ratios suggest that the reporter is suited to explore short-term ERK dynamics in ESCs.

We next recorded dynamic changes of reporter localization by imaging reporter cells at 20 second time intervals for up to two hours. In these time-lapse movies we could observe repetitive translocation of the sensor back and forth from the nucleus of cells growing in serum + LIF, which were absent in MEKi (Fig. 1F; Supp. Movie S1). To validate that these observations reflected genuine ERK activity, we transfected two spectrally compatible orthogonal ERK activity sensors in the same cells. Both sensors showed similar and highly correlated dynamic behavior (Fig. 1 Supp. 2). These sensors rely on different ERK substrate sequences, and deploy FRET (Komatsu et al., 2011) and translocation as two distinct read-outs. This indicates that pulsatile nuclear export of the KTR sensor reflects genuine ERK dynamics.

To quantify dynamic activity in single cells over time, we measured mean fluorescence intensity of the negative image in a region of interest within the nucleus (Methods). Thus, high values of the resulting KTR signal reflect high ERK activity, maintaining consistency with the representation in Fig. 1E. This analysis confirmed repeated pulses of sensor translocation in serum + LIF medium, which were suppressed by treatment with MEKi (Fig. 1G, Fig. 1 Supp. 3). We observed a broad range of dynamic behaviors across the population: some cells showed regular pulsing reminiscent of oscillations (* in Fig. 1G), and some showed isolated pulses (**). We also observed transitions between non-pulsing and pulsing behavior within the same cell (***). We conclude that ESCs display a range of pulsatile ERK activity dynamics when cultured in serum + LIF.

### Intermittent ERK oscillations in ESCs

The broad range of dynamic behaviors that we observed qualitatively across the population prompted us to systematically investigate the dynamic signatures of ERK activity in ESCs.

Since ERK activity pulses were a prominent feature of the dynamics, we sought to identify single pulses in time series. We first annotated the timepoints of local maxima and minima, and then used timeseries of MEKi treated cells to set a threshold for filtering ERK dependent pulses from background fluctuations (Fig. 2A, Fig. 2 Supp. 1, Supp. Table T1, Methods). Most cells (64/69, 93%) showed pulses in serum + LIF, while very few (2/67, 3%) showed any pulse in MEKi. The total fraction of time that single cells were pulsing was variable: some cells pulsed continuously, others showed a mixture of pulsing and non-pulsing behavior –termed silent– and yet others were non-pulsing throughout the experiment (Fig. 2B). On average, cells were pulsing (32 ± 3)% (mean ± SEM) of the time in serum + LIF alone, but only (0.13 ± 0.09)% of the time in the presence of MEKi (Fig. 2B).

**Fig. 2.**
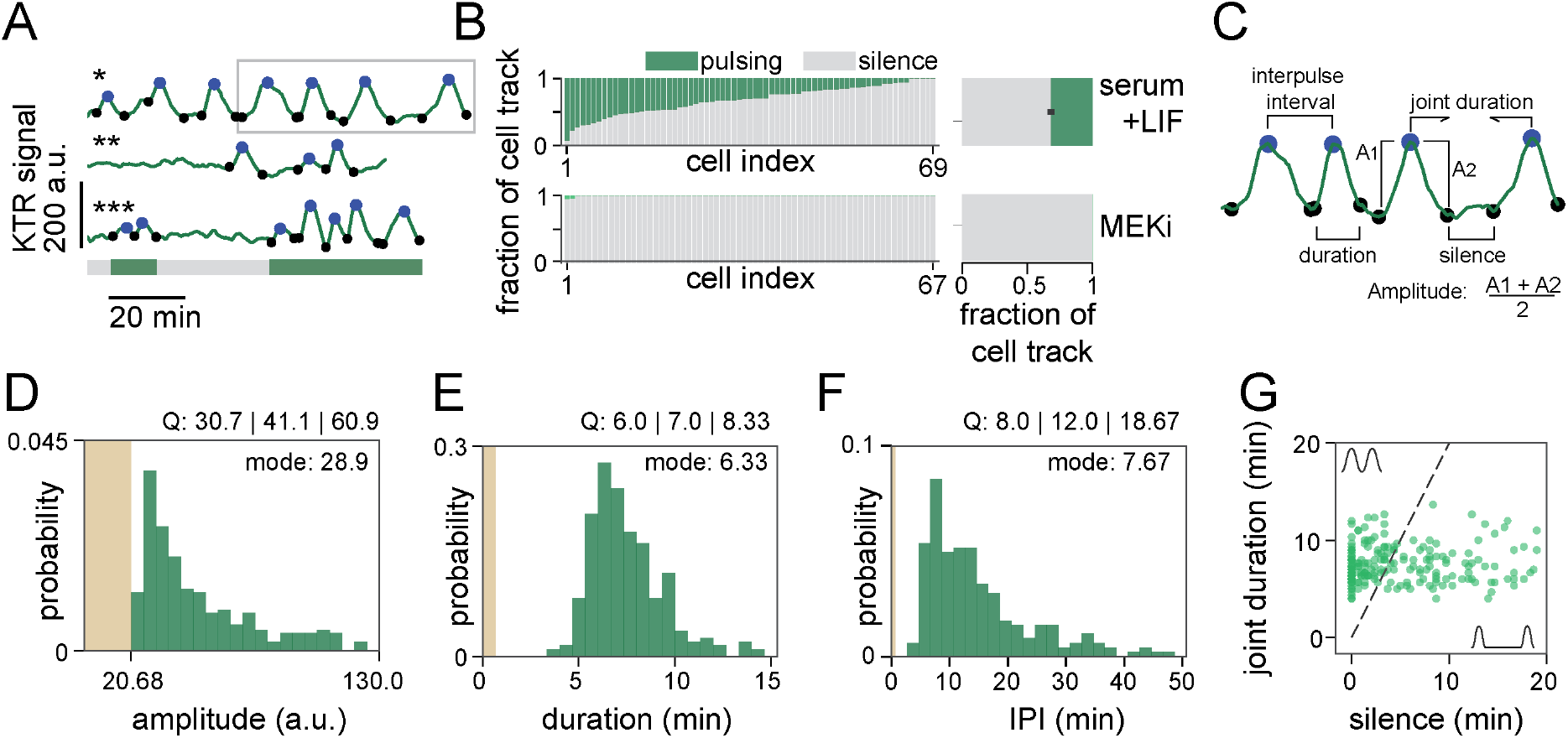
Time series analysis reveals intermittent ERK oscillations in ESCs. **A.** Pulse recognition in representative time series of ERK dynamical activity. Shown are smoothened single cell traces of KTR signal in the serum + LIF condition. Pulses are indicated by the maxima (blue dots) and corresponding minima (black dots) that define them. Bar at the bottom indicates pulsing (green) or non-pulsing (grey) intervals in the lower trace. **B. Left:** fraction of time that individual cells spent pulsing (green) or non-pulsing (grey) in serum + LIF alone (top) or upon addition of MEKi (bottom). **Right:** Average time that cells were pulsing (green) or non-pulsing (gray) in the cell population. Error bar indicates SEM**. C.** Dynamical features of the time series analyzed in **D-G** are indicated on a sample trace portion (gray rectangle in **A**). **D.** Pulse amplitude distribution for the serum + LIF condition (n = 289 pulses). **E.** Pulse duration distribution for the serum + LIF condition (n = 289 pulses). **F.** Interpulse interval distribution for the serum + LIF condition (n = 225 pairs of pulses). Pulse recognition resolution limit (yellow bar) and quartiles (Q) 25, 50 and 75 are indicated in **D-F**, and histograms are normalized to 1. **G.** Joint pulse duration vs. silence intervals for successive pairs of pulses in the serum + LIF condition (n = 225 pairs of pulses). The dashed line with slope 2 classifies pairs of pulses into consecutive (above) and non-consecutive (below). The axes range was adjusted to better resolve individual data points, leaving off the scale 27 out of 225 data points. Data in **D-G** from N = 69 cells.

To determine general characteristics of pulsing activity in the population, we introduced a set of quantitative measures: the amplitude and duration of single pulses, and the interpulse and silence intervals between successive pulses (Fig. 2C). The amplitude of a pulse was defined as the average difference between the peak value and the neighboring local minima (Fig. 2C). Our thresholding parameters only filter the tail of the amplitude distribution, containing low amplitude fluctuations that fall within the range of background levels determined from time series of MEKi-treated cells (yellow area in Fig. 2D).

We defined the duration of a pulse as the time elapsed between the two local minima flanking the maximum of the pulse (Fig. 2C, Methods). The distribution of pulse durations has a well-defined mode at 6.33 min and is slightly asymmetric (Fig. 2E). We observed no pulses shorter than 3 min, a timescale much longer than the detection limit of 40 seconds given by our algorithm and our sampling frequency. The congruence of the KTR and FRET sensors suggest that the pulse durations that we can capture are not limited by the timescales of sensor transport (Fig. 1 Supp. 2). Therefore, we conclude that ERK pulses have a minimum duration. Pulses with long durations tended to have large amplitudes, and those with short durations clustered at low amplitude values (Fig. 2 Supp. 2).

The interpulse interval (IPI) was defined as the time between the maxima of two neighboring pulses (Fig. 2C). The mode of the IPI distribution was 7.67 min, similar to the mode of the pulse duration (Fig. 2F). This suggests the presence of consecutive pulses, occurring immediately one after another. Consecutive pulses have either shared minima, or are separated by intervals of silence that are short relative to their pulse duration. As each interpulse interval can be decomposed into a silence interval and a joint pulse duration (Fig. 2C, Methods), we used these quantities to define consecutiveness in a way that accounts for differences in pulse duration. In a plot of joint duration against silence interval duration, sparse events will lie in the lower right region, while consecutive pulses will populate the upper left. Here we defined consecutive pairs of pulses as those with a silent interval of less than half the joint pulse duration (dashed line, Fig. 2G). With this definition, 52% of all pairs of pulses in cells growing in serum + LIF lay above the threshold and were classified as consecutive (Fig. 2G).

In summary, our analysis reveals that ERK pulses in ESCs growing in serum + LIF have a characteristic duration and are often part of consecutive sequences. We interpret this behavior as intermittent oscillations, where silent periods alternate with isolated pulses and oscillations – here defined as consecutive pulses with a characteristic duration.

### ERK oscillations are driven by FGF4

ERK activity is dynamic in many cell types (Albeck et al., 2013; Aoki et al., 2017; de la Cova et al., 2017; Goglia et al., 2020; Hiratsuka et al., 2015; Mayr et al., 2018; Pokrass et al., 2020; Shankaran et al., 2009; Simon et al., 2020). Extracellular signals can change the characteristics of these dynamics, such as pulse frequency (Albeck et al., 2013; Aoki et al., 2013). In ESCs, FGF4 is the main ligand that activates ERK (Kunath et al., 2007). We therefore asked how ERK dynamics depend on FGF4 concentration. To be able to control FGF4 concentration externally, we introduced an *Fgf4* loss of function mutation in the sensor line. In the chemically defined N2B27 medium that contains only minimal amounts of recombinant growth factors, *Fgf4* mutant cells were viable, but pERK levels were strongly reduced (Supp. Movie S2, Fig. 3 Supp. 1A). For stimulation, we chose FGF4 concentrations from 2.5 to 20 ng/ml. These concentrations cover the dynamic range of FGF4-response at the level of ERK phosphorylation (Fig. 3 Supp. 1B, C), transcription of an FGF/ERK-dependent reporter gene (Fig. 3 Supp. 1D, E), and differentiation along the primitive endoderm lineage (Raina et al., 2020). To measure the steady-state signalling response to different ligand levels, we pre-treated cells with the respective FGF4 concentrations for 24 h in pluripotency conditions, and replenished the medium 4 h before starting the recording (Fig. 3A, Methods). In the absence of FGF4 stimulation, we observed almost no pulsing. Widespread pulsatile activity was observed at all FGF4 concentrations tested, indicating that FGF4 triggers ERK pulsing (Fig. 3B, Fig. 3 Supp. 2, Supp. Movie S2). To identify pulses, we employed a similar strategy as above, setting a threshold based on the untreated condition and the highest FGF4 concentration (Fig. 3 Supp. 3, Methods).

**Fig. 3.**
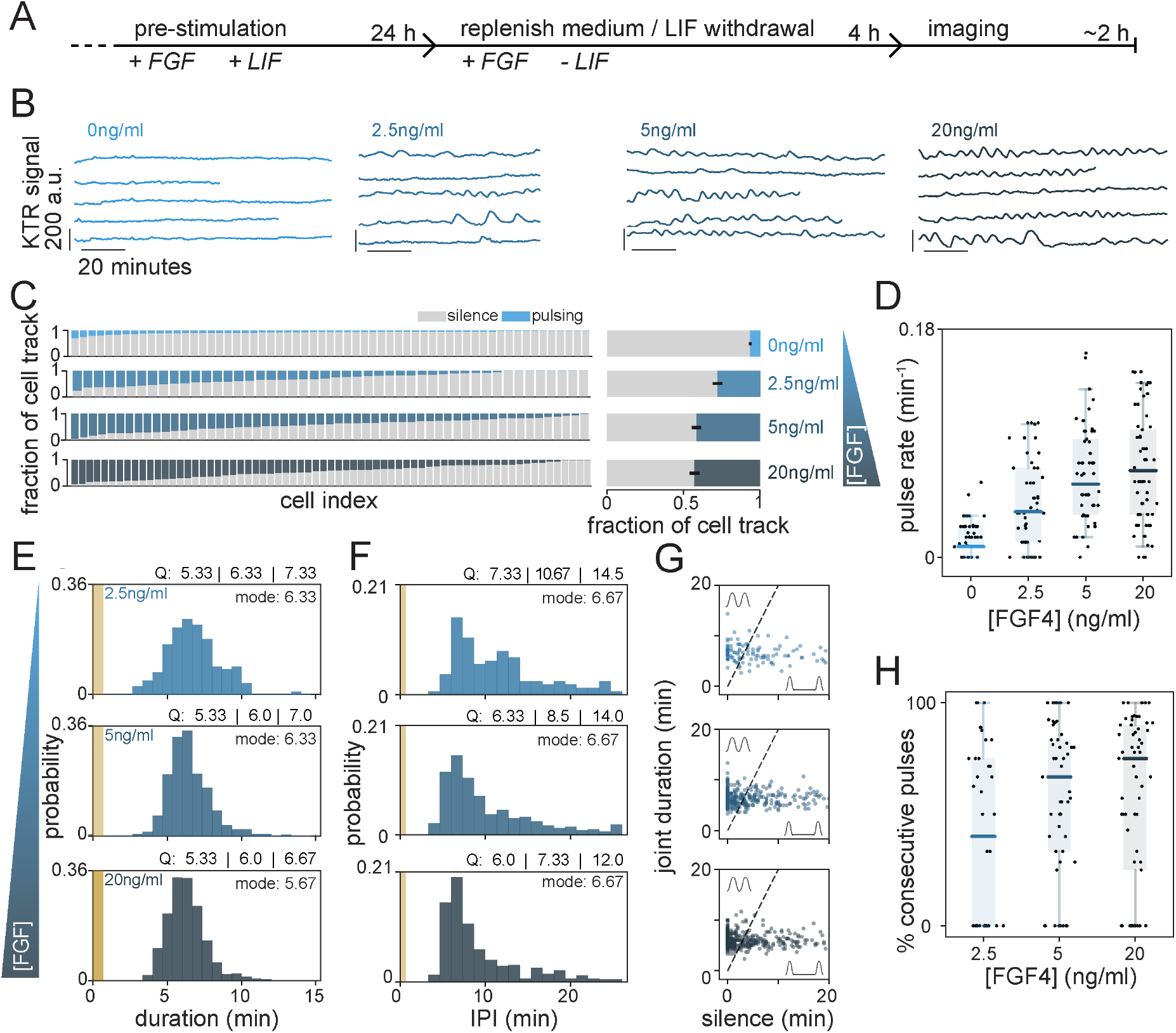
Pulsing and regularity of ERK activity are controlled by FGF4 dose. **A.** Schematic of experimental protocol to measure the steady state signaling response to defined FGF4 ligand levels. **B.** Representative traces of smoothened time series of ERK dynamical activity in single Fgf4 mutant cells stimulated with different FGF4 doses. **C. Left:** Fraction of time that individual cells stimulated with different concentrations of FGF4 spent pulsing (blue), or non-pulsing (grey). **Right:** Average time that cells in the population were pulsing (blue) or non-pulsing (gray). Error bar indicates SEM**. D.** Pulse rate boxplots at different concentrations of FGF4. **E.** Pulse duration distributions. The number of pulses was n = 164 (2.5ng/ml), n = 426 (5ng/ml) and n = 544 (20ng/ml). **F.** Distributions of interpulse intervals between pairs of successive pulses. The number of successive pulses was n = 124 (2.5ng/ml), n = 370 (5ng/ml) and n = 479 (20ng/ml). Pulse recognition resolution limit (yellow bar) and quartiles (Q) 25, 50 and 75 are indicated in **E** and **F**, and histograms are normalized to 1. **G.** Joint pulse duration vs. silence intervals for successive pairs of pulses. The slope 2 dashed line classifies pairs of pulses into consecutive (above) and non-consecutive (below). The axes range was adjusted to better resolve individual data points, leaving off the scale 6 of 124 (2.5ng/ml FGF4), 26 out of 370 (5ng/ml FGF4) and 33 out of 479 (20ng/ml FGF4) data points. Number of cells in **C**–**G**: N = 61 (0ng/ml FGF4), N = 48 (2.5ng/ml FGF4), N = 57 (5ng/ml FGF4) and N = 69 (20ng/ml FGF4). **H.** Ratio of consecutive pulses to the total number of pulses in single cells. Number of cells was N = 41 (2.5 ng/ml), N = 56 (5 ng/ml) and N = 67 (20 ng/ml), cells with no pulses were not included. Box plots (**D**, **H**): Black dots represent individual cells, color bars are the median, box bounds are the 25 and 75 percentiles of the distributions, and whiskers are the 5 and 95 percentiles.

The distribution of sensor pulse amplitudes was not significantly different amongst the three concentrations (Fig. 3 Supp. 4A, Methods, Supp. Table T2). However, immunostaining and single cell analysis revealed that the median, the lower end, as well as the variance of the pERK distributions shifted to larger values with increasing FGF concentration (Fig. 3 Supp. 4B, C). Thus, it is possible that the amplitude of pERK pulses increases with FGF concentration, without translating into a measurable increase in sensor pulse amplitude.

The total fraction of time that single cells were pulsing increased with FGF4 concentration in the range from 0 to 5 ng/ml (Fig. 3C), to levels similar to those measured in wild type cells in serum + LIF. We wondered how the number of pulses and their duration contributed to this increase in pulsing time. We defined a single cell pulse rate as the number of pulses divided by the duration of the trace, and found that it increases with FGF4 concentration in the same range (Fig. 3D). The distribution of pulse durations overlapped between the three FGF4 concentrations, and their modal values were conserved (Fig. 3E). We observed a subtle trend towards narrower distributions with higher FGF4 concentrations, yet these were significantly different only between the 2.5 ng/ml and 20 ng/ml conditions (Supp. Table T2, Methods). Thus, the increase in pulsing time is largely due to an increase in pulse rate rather than pulse duration. In line with stable pulse durations, the IPI distributions had a similar modal value of about 7 min in all conditions. However, IPIs became more narrowly distributed with increasing FGF4, with a clear difference between 2.5 and 20 ng/ml FGF4 (Fig. 3F, Supp. Table T2). Narrower IPI distributions at high FGF concentrations indicated more regular pulsing.

To determine whether the extent of consecutive pulsing was controlled by FGF4, we plotted joint pulse duration against silence interval duration (Fig. 3G). In the population, the fraction of consecutive pulses increased steadily across the entire FGF concentration range from 49.2% (2.5 ng/ml) to 57.6% (5 ng/ml), and 63.9% (20 ng/ml). We counted isolated and consecutive pulses in single cell traces to evaluate their contribution to this population behavior. Here, isolated pulses include those from non-consecutive pairs as well as those from traces with single pulses in which no silence intervals were defined. The proportion of cells showing only isolated pulses decreased from 43% (17 out of 40) at 2.5 ng/ml FGF4 to 18% (10 out of 56) and 24% (16 out of 65) at 5 ng/ml and 20 ng/ml, respectively. In the cells that showed consecutive pulsing, the fraction of consecutive pulses increased with FGF4 across the entire concentration range that we tested (Fig. 3H).

In summary, these results reveal that ERK pulses have a characteristic duration that is independent from FGF4 concentration. The increase of both pulse rate and consecutiveness, together with the narrowing of duration distributions, suggest that FGF dose controls the extent as well as the precision of ERK oscillations.

### ERK pulses are more prevalent early in the cell cycle

We noted that within the same experimental condition, there was significant cell-to-cell variability in pulsing activity (Fig. 2B, Fig. 3C). This observation could result from stable differences in pulsing behavior between cells. Alternatively, single cells could transition back and forth between pulsing and non-pulsing states, which would show up as different behaviors when observation times are limited in comparison to the characteristic times of such transitions. To identify changes in pulsing behavior of single cells over longer timescales, we recorded movies for 18 hours such that cells could be followed from birth to division (Fig. 4A, Fig. 4 Supp. 1). Increasing the frame intervals to 105 s reduced overall light exposure, while still allowing to resolve pulses that are at least 3.5 min apart. We recorded pulsing in wild type cells growing in N2B27 medium, thereby exclusively focusing on pulsing driven by paracrine FGF4 signaling, and avoiding possible ligand depletion that could occur with exogenous FGF4. We established an alternative peak-finding approach to quantify and annotate these low temporal resolution traces (Fig. 4B, Fig. 4 Supp. 2, Methods). We made raster plots showing occurrence of pulses in cells that we could follow from immediately after cell division (Fig. 4C). Visual inspection of these raster plots suggested that pulses were concentrated towards the beginning of the cell cycle. A change in pulsing activity over time could be a consequence of cell cycle effects on pulsing, or it could result from non-stationary experimental conditions.

**Fig. 4.**
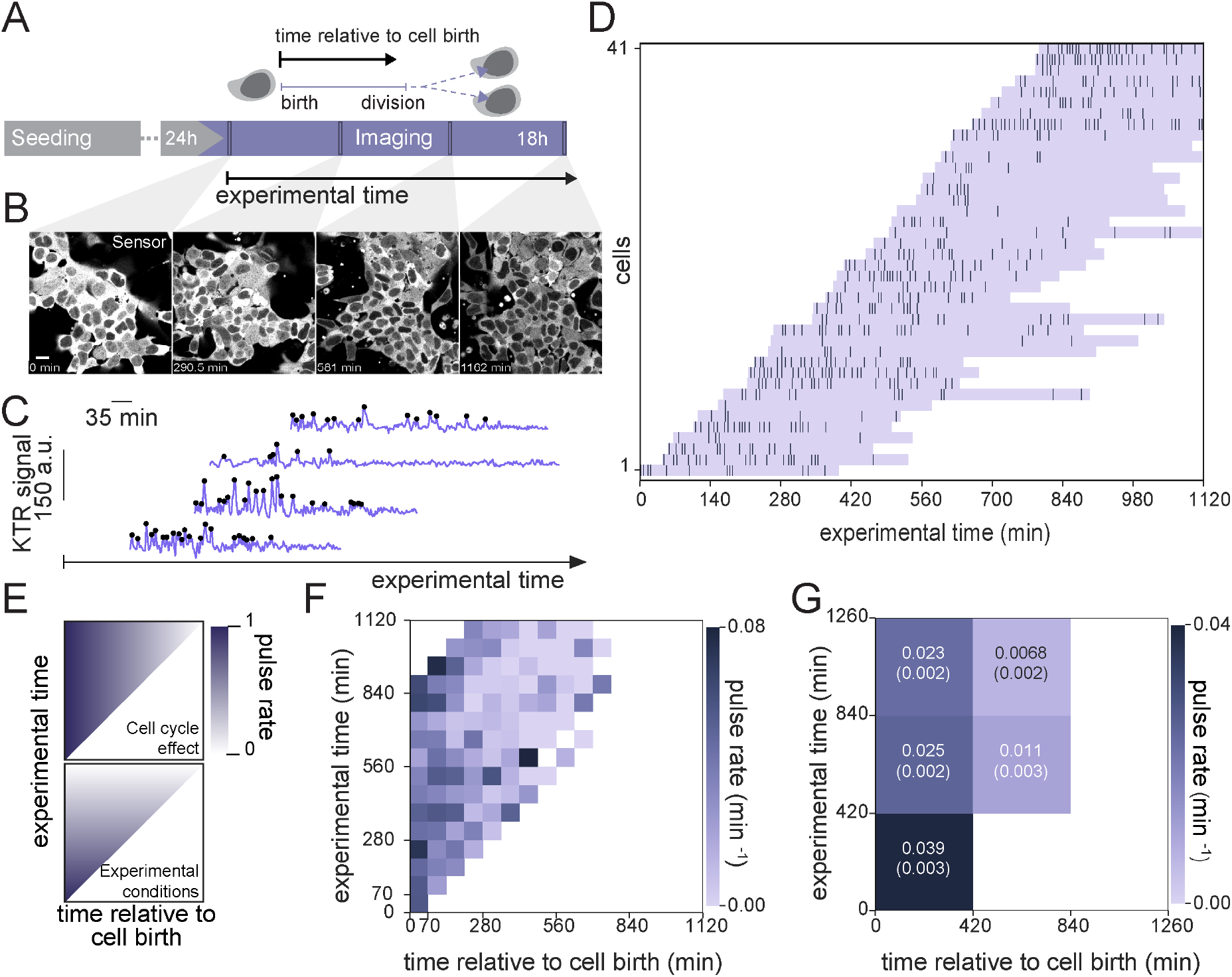
ERK pulsing is more prevalent early in the cell cycle. **A.** Schematic of experimental protocol to record ERK peaks across complete cell cycles. **B.** Montage of an ESC colony expressing the ERK-KTR sensor over the course of a long-term imaging experiment. Scale bar = 20 μm. **C.** Representative filtered traces of ERK dynamical activity with identified peaks (black dots), in single wild type cells growing in N2B27 medium. **D.** Raster plot displaying the timing of ERK activity peaks across the cell cycle. Lavender horizontal bands extend from birth to division of single cells, dark vertical bars represent peaks. Single cell tracks begin immediately after a cell division event and are plotted relative to absolute experimental time. **E.** Schematic representation of expectations for a reduction of pulsing activity due to cell cycle (top) and due to changing experimental conditions (bottom) in the 2-dimensional color-encoded pulse rate map. **F.** Pulse rate map for the data shown in **D**. Time is discretized into 70 min bins. **G.** Coarse grained pulse rate map showing average pulse rate and its estimated error with 420 min binning.

To visualize the contributions from these two possible causes, we introduced a two-dimensional time map. The coordinates in this map are experimental time *T_e_*, which is time measured from the beginning of the time lapse movie and *T_b_*, the time relative to individual cell birth. For each cell *i,* the trace begins at 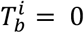 and experimental time 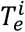, the time when cell *i* was born measured from the beginning of the movie. From this point in the map, individual traces would fall along a diagonal line of unit slope. To reveal the population behavior and avoid superposition of individual traces in the map, we plot pulse rate averaged in 70 min bins along both axes. In each bin, we count the total number of pulses from all traces in that bin and divide by the total number of minutes of recording that contribute to that bin. On this pulse rate map, cell cycle effects would manifest as a rate change in the horizontal direction (Fig. 4E, upper panel) while non-stationary experimental conditions would manifest as a change in rate in the vertical direction (Fig. 4E, lower panel).

Inspection of the pulse rate time map revealed a higher pulse rate at the bottom left of the plot that decreased both towards the right and the top (Fig. 4F). This behavior indicates that pulse rate decays across the cell cycle, in addition to effects of non-stationary experimental conditions. To quantify this observation, we further binned pulse rate at larger timescales (Fig. 4G). In this coarse-grained map, pulse rate within the same experimental time window was consistently higher in cell populations which were earlier in their cell cycles. We obtained similar results when applying an alternative detrending strategy (Fig. 4 Supp. 2, Fig. 4 Supp. 3), as well as when analyzing cells growing in serum + LIF medium (Fig. 4 Supp. 4). Taken together, these results confirm that cells are more prone to pulse earlier in their cell cycle.

## Discussion

Here we report fast pulses of ERK activity in mouse embryonic stem cells under a continuous stimulation regime. We demonstrate that this pulsing activity is consistent with oscillations, with transitions between silent and oscillatory states in single cells. Oscillations are driven by FGF4. Across a range of FGF4 ligand concentrations, we find oscillations with similar individual pulse durations. With increasing FGF4 concentrations, the distribution of interpulse intervals becomes narrower and the fraction of consecutive pulses increases, suggesting more regular oscillations.

The detection of signal-dependent ERK activity dynamics on short time scales in ESCs was made possible by combining the KTR sensor with high time-resolution recordings. A previous study which examined ERK dynamics upon acute stimulation focused on long term activity and did not resolve the short-timescale oscillations that we report here (Deathridge et al., 2019). These previously undetected dynamics have a modal interpulse interval of approximately 7 minutes (that is, about 8 pulses per hour), and are thus much faster than in any other cell system described so far.

Both paracrine and recombinant FGF4 stimulation of ESCs trigger oscillatory ERK activity with similar timescales of pulse duration and IPI, indicating that oscillations emerge in the intracellular signal transduction network, similar to the situation in other cell lines (Sparta et al., 2015). The short frequencies of ERK oscillations in ESCs further support the notion that they are driven by short-timescale delayed feedbacks such as post-translational modifications at the receptor level (Sparta et al., 2015), or at various levels within the MAPK cascade (Lake et al., 2016; Lemmon et al., 2016).

Pulsatile ERK activity in single cells upon continuous stimulation of RTKs has been reported in many cell types (Albeck et al., 2013; Goglia et al., 2020; Shankaran et al., 2009), indicating that the tendency to generate time-varying ERK activity patterns is a general feature of RTK signal transduction. In addition to the timescales, the dynamic signatures of FGF-triggered ERK pulses in ESCs however differ markedly from those observed in most other contexts.

ERK pulses in ESCs have well-defined durations and narrowly distributed IPIs, consistent with oscillations. This is in contrast to the more irregular, stochastic pulsing reported in several immortalized cell lines and keratinocytes (Albeck et al., 2013; Aoki et al., 2013; Goglia et al., 2020). Regular oscillations of ERK nuclear import and export have been reported in mammary epithelial cells (Shankaran et al., 2009). In these cells, the frequency of ERK oscillations is insensitive to ligand levels over a wide range (Shankaran et al., 2009), similar to what we find upon titrating FGF4 in ESCs. Remarkably, across a wide range of ligand levels ESC populations contain a mixture of oscillating and non-oscillating cells as well as cells that transition between these regimes. This suggests that the FGF/ERK signal transduction system in ESCs is organized in the vicinity of a transition point between a non-oscillatory and oscillatory state. In this framework, increasing FGF4 levels would bring the system closer to this point. Similarly, the decay of ERK pulsing across the cell cycle can be interpreted as cells shifting away from the oscillatory to a non-oscillatory state, possibly through changes in the surface-to-volume ratio or cell cycle-dependent expression of components of the FGF/ERK signaling system. Such positioning close to a transition between oscillatory and non-oscillatory behavior has been described in hair cells of the cochlea (Camalet et al., 2000; Eguíluz et al., 2000), the actin system of *Dictyostelium* (Westendorf et al., 2013), and isolated cells of the growing vertebrate body axis (Webb et al., 2016), suggesting that this is a generic principle. The mechanism that positions the FGF/ERK signaling system in ESCs close to this transition point, the molecules involved, and the possible physiological relevance of being close to this transition, remain to be identified.

In cell systems that show stochastic ERK pulsing, increasing ligand levels leads to shorter interpulse intervals and hence to an increase in pulse rate (Albeck et al., 2013). This has been interpreted as frequency-modulated (FM) encoding of ligand concentration. In ESCs, the mode of the interpulse intervals is largely independent from FGF4 concentration, and pulse rate in the population increases mostly as a consequence of an increase in the fraction of time that individual cells spend pulsing. Thus, the FM-encoding model proposed for stochastic ERK pulsing is unlikely to apply in differentiating ESCs.

Still, the signalling dynamics that we report here reflect the cell type-specific organization of the FGF/ERK system in ESCs. While single cell models for FM-encoding are based on excitable dynamics arising from a combination of positive and negative feedback loops (Aoki et al., 2013; Tsai et al., 2008), oscillations with regular interpulse intervals require delayed negative feedback only (Novák and Tyson, 2008). Frequency-modulated ERK pulses are often found downstream of the EGF receptor, for which autocatalysis provides a positive feedback mechanism (Koseska and Bastiaens, 2020). Positive feedback might be less prominent for the FGF receptor in ESCs, such that ERK dynamics is dominantly shaped by negative feedback. Negative feedback in the RAF-MEK-ERK cascade sets the ligand dose response range, and linearizes signal transduction despite non-linear signal amplification (Sturm et al., 2010). In ESCs and cells of the early embryo, the proportion of differentiated cell types smoothly depends on FGF4 concentration (Krawchuk et al., 2013; Raina et al., 2020). The oscillatory ERK activity that we detect here might be a consequence of negative feedback mechanisms that have evolved to tune the response range of the signal transduction system to the physiologically relevant range of paracrine FGF4 concentration, and faithfully transmit this information to the transcriptional level. Interfering with candidate mechanisms for negative feedback will be required to establish the connections between network architecture, oscillations, and cell differentiation.

Our identification of heterogeneous signaling dynamics adds another dimension to the phenomenon of cellular heterogeneity which is a hallmark of embryonic stem cell cultures in vitro (Canham et al., 2010; Chambers et al., 2007; Hayashi et al., 2008; Singh et al., 2007; Toyooka et al., 2008). Consistent with the well-known increase in cellular heterogeneity in serum + LIF (Kalkan and Smith, 2014), we observe a broader distribution of IPI in this culture condition compared to defined conditions. Heterogeneities in stem cell cultures have classically been attributed to the noisy activity of gene regulatory networks that control cell state. Correlating signaling dynamics with the state of transcriptional networks over time will be required to discern how signaling heterogeneities are causally related to these transcriptional cell states.

## Acknowledgements

We thank Michelle Protzek and Holger Vogel for technical support, and Philippe Bastiaens and Heinz Neumann for suggestions and input at early stages of the project. We thank Aneta Koseska, Koichiro Uriu, John Albeck, Alfonso Martinez Arias and members of our groups for helpful suggestions on earlier versions of this manuscript. FF acknowledges funding from CONICET and DAAD, and the hospitality of the Department of Systemic Cell Biology. LM acknowledges funding from ANPCyT PICT 2017 3753. Work in our groups is supported by FOCEM-Mercosur (COF 03/11) and the Max Planck Society.

## Methods

### Cell culture

mESCs were routinely cultured on 0.1% gelatin (Sigma Aldrich)-coated tissue culture flasks in serum + LIF medium composed of GMEM (ThermoFisher), 10% batch-tested fetal bovine serum (FBS) (Sigma Aldrich), 1x GlutaMAX (ThermoFisher), 1 mM sodium pyruvate (ThermoFisher), 1x non-essential amino acids solution (ThermoFisher), 100 μM 2-mercaptoethanol (ThermoFisher) and 10 ng/ml LIF (MPI protein expression facility). Cells were passaged every two to three days using 0.05% Trypsin (PAN Biotech). Basal medium for serum free culture was N2B27, prepared as a 1:1 mixture of DMEM/F12 (PAN Biotech) and Neuropan basal medium (PAN Biotech) with 0.5% BSA, 1x N2 and 1x B27 supplements (ThermoFisher) and 50 μM 2-mercaptoethanol. For FGF stimulation experiments, short-term serum-free culture was carried out in N2B27 supplemented with 3 μM CHIR99201 (Tocris), 1 μg/ml of Heparin (Sigma) and with or without 10 ng/ml LIF as indicated. Recombinant human FGF4 used was obtained from Peprotech. For live imaging and immunostaining studies, cells were seeded on polymer-bottomed ibidi μ-slides (ibidi) coated with 20 μg/ml fibronectin.

### Cell lines

All KTR-expressing cell lines used in this study were derived from E14tg2a (Hooper et al., 1987). Targeting of the ERK-KTR-Clover construct into the *Hprt* locus has been described elsewhere (Simon et al., 2020). Mutagenesis of the *Fgf4* gene was performed by cotransfection of a CRISPR-construct and a repair template introducing a nonsense and a frameshift mutation as previously described (Morgani et al., 2018). Clones with the desired mutation were identified by restriction digest and Sanger sequencing of a PCR fragment encompassing the *Fgf4* start codon. Clonal cell lines were tested for their chromosome count using standard procedures (Nagy et al., 2008), and only cell lines with a modal count of n = 40 were used for analysis. *Fgf4^-/-^, Spry4^H2B-Venus/H2B-Venus^* cells to evaluate transcriptional activation downstream of recombinant FGF4 have been described (Morgani et al., 2018).

### Dual reporter experiments

The ERK-KTR-mCherry construct for transient expression was prepared by first inserting the coding sequence for ERK-KTR (Regot et al., 2014) into a CMV-driven mCitrine C1 expression vector (TaKaRa), and then replacing the fluorophore for mCherry. The plasmid for transient expression of EKAREV-NLS has been described (Komatsu et al., 2011). 1.5 μg each of plasmid were transiently transfected into E14tg2A mouse stem cells using Lipofectamine 2000 (ThermoFisher) in suspension according to manufacturer’s instructions. Cells were plated on fibronectin-coated ibidi slides and imaged 24h after transfection.

### Western blotting

Cells were grown to confluency on fibronectin-coated tissue culture dishes and exposed to indicated experimental conditions. Cells were briefly washed twice with ice-cold PBS supplemented with 1 mM activated sodium orthovanadate and then lysed using commercially available lysis buffer (Cell Signaling) supplemented with benzonase (ThermoFisher), phosphatase inhibitor cocktail 2 and 3 (Sigma), and cOmplete EDTA-free protease inhibitor cocktail (Roche). Lysates were snap-frozen in liquid nitrogen. Protein concentration was estimated using a micro-BCA assay (ThermoFisher), and lysates were denatured by adding appropriate amounts of 5x Laemelli buffer and boiling for 5 min. 10 or 20 μg protein was loaded across all wells in any given gel. Bis-Tris SDS gels were run with 1x MOPS buffer (ThermoFisher) with fresh sodium bisulfite, and subsequently transferred onto methanol-activated PVDF membranes (Millipore) at 40 V for 1.5 h with the NuPage transfer system (ThermoFisher). Primary antibodies used were anti-Tubulin 1:5000 (T6074, Sigma), anti-pERK1/2 1:1000 (4370S, Cell Signaling), and anti-total ERK1/2 1:1000 (ab36991, Abcam) along with appropriate secondary antibodies (LI-COR). Bands were detected using the Odyssey CLx imaging system (LI-COR). Bands were quantified using FIJI/ImageJ (Rueden et al., 2017). For quantification of pERK and total ERK, integrated intensity in both ERK1 and ERK2 bands was added.

### Immunostaining

For pERK immunostaining, cells were fixed for 15 min at 37°C by diluting fixative stocks directly into cell culture medium to a final concentration of 4% PFA and 0.01% glutaraldehyde (Sigma). After a brief wash with PBS, cells were permeabilized with 100% methanol at −20°C. For all other antibodies, fixation was performed with 4% PFA at room temperature for 20 min. Cells were washed with PBS and then simultaneously blocked and permeabilized with 5% normal goat serum (ThermoFisher) in 0.5% Triton X-100 (Serva) in PBS for 60 min. Antibody staining was carried out overnight at 4°C in PBS + 0.1% Triton X-100 and 1% BSA (Sigma). Primary antibodies used were anti-pERK1/2 1:200 (4370S, Cell Signaling), anti-E-Cad 1:200 (M108, clone ECCD-2, TaKaRa), anti-Nanog 1:200 (eBIO-MLC51), anti-POU5F1 1:200 (C10, sc-5279, Santa Cruz), along with appropriate secondary antibodies. Hoechst 33342 was used at 1μg/ml to counter-stain nuclei, and CellMaskRed (ThermoFisher) was used to label membranes according to manufacturer’s instructions. After staining, samples were covered with 200 μl of antifade composed of 80% w/v glycerol with 4% w/v N-propyl gallate and stored at 4°C. Images were analyzed using custom scripts in MATLAB (The Mathworks) and Fiji/ImageJ for the detection of nuclei as well as an active-contours based identification of membranes.

### Flow cytometry

Cells were grown on fibronectin-coated dishes in N2B27 supplemented with 3 μM CHIR99201, 1 μM PD0325901 and 10 ng/ml LIF (2i + LIF) for 3 days. For stimulation, cells were washed 2x with PBS, and FGF4 was added at indicated concentrations in serum-free N2B27 medium supplemented with 3 μM CHIR99201 and 1 mg/ml Heparin for 24 h. Cells were then trypsinized and fixation was performed in suspension with 4% paraformaldehyde at room temperature for 15 min. After a brief wash in PBS, cells were resuspended in PBS + 1% BSA and analyzed on a BD-LSR II (BD Biosciences) flow cytometer. Data was analyzed in FlowJo (BD Biosciences).

### Live cell imaging and cell tracking

ERK-KTR expressing cells were cultured on ibidi μ-slides, and imaged on a Leica SP8 confocal microscope equipped with an incubation chamber and CO2 supply to maintain temperature at 37°C, CO2 at 5%, and relative humidity at 80%. 4 h before acquisition, live-cell nuclear dye SiR-Hoechst 652/674 (Spirochrome) was added to facilitate tracking of cells. SiR-Hoechst was added at a final concentration of 500 nM for short-term time-lapse experiments, and 250nM for long-term time-lapse experiments. Fluorophores were excited with a 504nm line from a white-light laser (Leica), and images of the KTR-Clover and the nuclear marker were simultaneously captured through a 63x 1.4 N.A. oil objective. For short-term (~2 h) imaging experiments, single frames were acquired once every 20 s, with an XY resolution of 0.251 nm, with a pixel dwell time of 2.6 μs, and a pinhole of 2.4 airy units. For long term (~19 h) imaging experiments, to minimize overall light exposure single frames were acquired once every 105 s, with an XY resolution of 0.401 nm, with a pixel dwell time of 3.1 μs, and a pinhole of 2.6 airy units. Images were processed with custom MATLAB scripts to enhance contrast and highlight nuclei to facilitate automatic tracking. Tracking was performed using the Trackmate plugin (Tinevez et al., 2017) for FIJI/ImageJ. Tracking was initially performed automatically for the entire colony, and tracks were subsequently manually curated frame-by-frame by removing any cells that did not display a typical ESC morphology with a small cytoplasm and round, well-defined nuclei. We also removed cells that left the field of view, and adjusted tracking in individual frames for incorrectly identified nuclei. We inverted fluorescence values to obtain the negative image, and then measured mean fluorescence intensities in a region of interest (ROI) of variable size within each tracked nucleus. In these KTR signal traces, low intensity values correspond to low ERK activity and high intensity values indicate high ERK activity. For the short-term imaging, tracks started at the beginning of the movie and extended until the end of the movie, or until cell division. As the long-term imaging experiments were designed to capture the entire cell cycle, tracks started in the first frame following cell division where a cell could be tracked, and ended at cell division. In these experiments, we kept tracks of cells that left the field of view, but only if they were observed for longer than 4.5 h.

### Time series preprocessing

We screened and corrected time series for tracking errors, such as ROIs placed partially outside the nucleus or overlapping with a nucleolus. Because these structures have fluorescence intensities that usually differ from that of the nucleoplasm, these tracking errors usually led to an increase in the variance of the pixel intensity across the ROI. We screened time series for high variance regions, checked the tracking for all instances where the variance crossed a manually set threshold value, and corrected the tracking if this was required.

Just before cell division, the sensor was excluded from the nucleus, resulting in a pulse of the KTR signal at the end of dividing cells tracks (for example cells 30, 31, 41 and 50 in the serum + LIF condition without MEKi (Fig. 1 Supp. 2)). As this pulse of reporter exclusion was insensitive to MEK inhibition, it is unlikely reporting ERK activity and we therefore decided to trim these events from all traces. While most cells divided in the long-term measurements, only a few did it in short-term measurements. Correspondingly, in short-term measurements we deleted the last 20 frames (about 7 min) of the time series of dividing cells only. In the long-term measurements, where most cells divided, we discarded the last 15 frames (20.25 min) of each time series.

### Analysis of ERK dynamics in short-term high resolution datasets

#### Pulse recognition

We defined a pulse as a local maximum between two local minima, imposing two conditions: (i) we required amplitude to be larger than a threshold amplitude *A_th_,* and (ii) slope to be larger than a threshold slope *v_th_.* The amplitude and slope thresholds are free parameters of the algorithm. These free parameters were set through a quantitative threshold analysis protocol described below and were specific for each dataset (Supp. Table T1).

To remove high frequency noise that interfered with the performance of the pulse detection algorithm, we first smoothed the time series (Fig. 2 Supp. 1, Fig. 3 Supp. 3). We filtered the highest frequencies in the data using a moving average window of 3 frames of duration. That is, for each KTR signal value *x_i_* of the time series, we computed the average value

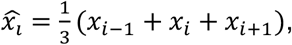

where *i.* is the frame number. At the boundaries we considered average windows of 2 and 1 frames. Note that detrending was not required in the case of this data.

We first searched the time series for all the local maxima and minima. We compared each value 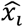 of the time series with its immediate neighbours 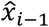 and 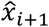. The initial value 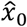 was compared only with the next value 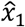 and the last value 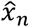 with the previous one 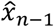 (Fig. 2 Supp. 1, Fig. 3 Supp. 3). We discarded the first maximum if there was no minimum on its left, and the last maximum if there was no minimum on its right. In this way we defined a subset of data points consisting of the maxima 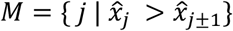, and the subset of minima 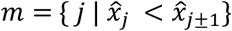. From the definition, it follows that the minimum distance |*i* – *j*| between two maxima 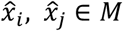 is 2 frames, and the minimum distance |*k* – *l*| between two minima 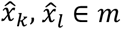 is 2 frames.

To identify pulses from this set of maxima we applied two filters, one for pulse amplitude and another one for pulse slope. To implement the pulse amplitude and pulse slope filters we considered each maximum of the time series, from left to right. For each maximum *j* ∈ *M* of 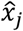, we searched for the first minimum to its left *k* ∈ *m* such that the resulting left amplitude 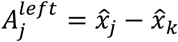 was larger than the amplitude threshold 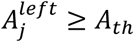 and the left slope was larger than the slope threshold 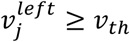 (see threshold analysis protocol below). The left slope was defined as 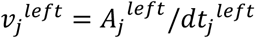, where 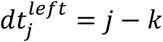 is the left pulse duration. Similarly, we searched the first minimum to the right that verified 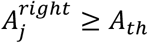 and 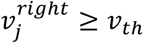. We next removed overlapping pulse candidates: if the right minimum of the first pulse occurred later than this new left minimum of the second one, we discarded the pulse that had the smaller amplitude 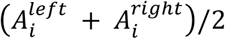 (Fig. 2 Supp. 1, Fig. 3 Supp. 3).

#### Threshold analysis protocol

Pulse recognition depends on the free parameters for amplitude threshold *A_th_* and slope threshold *v_th_.* To rationally set values for these two threshold parameters, we first focused on the negative control condition for each respective experiment, where ERK pulsing was minimal. We determined parameter combinations for which a fixed, low number of pulses was detected in the negative control, and then selected specific parameter values that maximized the number of pulses recognized in the experimental condition where ERK was most active (Supp. Table T1).

We started by establishing a two-dimensional exploratory parameter space for each dataset (Supp. Table T1). For each combination of parameters 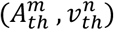 on the exploratory parameter space, we run the pulse detection algorithm described in the previous section for the negative control and computed the averaged pulse rate

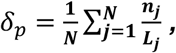

where *N* is the total number of cells in the negative control, *n_i_* is the number of detected pulses for cell *j* and *L_j_* is the length of the time series (Fig. 2 Supp. 1). We then introduced exploratory level curves across the parameter space by fixing average pulse rate values 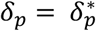 in the negative control (Supp. Table T1). This restricted parameter combinations to curves in the exploratory parameter space (Fig. 2 Supp. 1, Fig. 3 Supp. 3). Next, for each 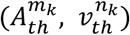 combination on each exploratory level curve *k,* we applied the pulse recognition algorithm on the experimental condition where ERK was most active. The plot of pulse rate along this level curve showed a flat region of similarly high pulse detection. Within this region, we chose parameters pairs that filtered out spurious pulses that were flat and long from the negative control. This resulted in a pair of parameters (*A_th_, v_th_*) specific for each experiment (Fig. 2 Supp. 1, Fig. 3 Supp. 3, Supp. Table T1).

#### Quantitative pulse dynamics characterization

To characterize dynamical activity of the time series, we introduced a set of quantitative measures (Fig. 2D). For each pulse *P$* in the set of pulses *P* = {*P_j_* > = (*j, k_j_, l_j_*) | *P_j_ is a pulse* } we defined the **pulse amplitude** *A_i_* as the average of its right and left amplitudes

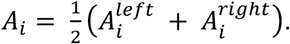

**Pulse duration** *dt_i_* was defined as the distance between the two minima that define the pulse

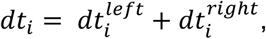

and the **joint pulse duration** *dt_ij_* between a pair of successive pulses *P_i_, P_j_* with *j* > *i,* as the sum of the right pulse duration of the earlier pulse *i.* and the left pulse duration of the later pulse *j*

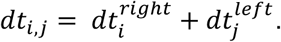

We computed the **interpulse interval** *IPII_i,j_* between a pair of successive pulses *P_i_, P_j_* with *j* > *i,* as the time interval between their maxima

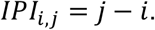

The **silent interval** *dm_i,j_* between a pair of successive pulses *P_i_, P_j_* was defined as the time elapsed between the right minimum of the earlier pulse *P_i_* and the left minimum of the later pulse *P_j_*, that is

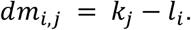

Note that calculating these last three quantities requires a trace with at least two pulses. These quantities satisfy the relationship

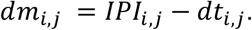

The values that these quantitative measures can take are constrained by the resolution imposed by pulse recognition. The minimum distance |*i* – *j*| between two maxima 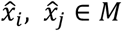 was previously set to 2 frames. Thus, the distance between maxima of pulses *P_k_, P_l_ ∈ P* verifies |*k* – *l*|≥2 frames, and in particular *IPI_k,l_* ≥ 2 frames for any pair of consecutive pulses *P_k_,P_l_* ∈ *P*. Similarly, the minimum distance |*i* – *j*| between two minima 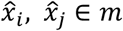 is 2 frames. Consequently, given a pulse *P_j_* = (*j, k_j_, l_j_*) ∈ *P*, the distance between the two minima that defines the pulse *dm_j_* = *k_j_* – *l_j_* satisfies *dm_j_* ≥ 2 frames. Finally, from the previous section we have the constraints *A_i_* > *A_th_* and *v_i_* > *v_th_*.

We classified pulses as **consecutive** or **isolated**. Inspection of the raw data indicated that pulse duration was more variable between cells in the same condition than within a cell. For this reason, we made the criterion for consecutiveness dependent on joint duration of the half-pulses that flank an intervening silent period. Specifically, we established that a pair of successive pulses *P_i_, P_j_* are **consecutive pulses** if the silent interval between them *dm_i,j_* is shorter than half of their joint duration *dt_i,j_*, that is *P_i_, P_j_* are consecutive if *dm_i,j_* ≤ 0.5 *dt_i,j_*. Pulses that do not belong to a consecutive pair are **isolated pulses**.

We also introduced a quantitative measure to characterize the dynamical activity on a population level. Given a single cell *c* associated to a time series of total length *T* and *n* pulses, the **pulsing** measure *A_c_* is defined as the proportion of time that a single cell is pulsing

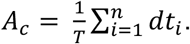

#### Kolmogorov-Smirnov test and notation

We implemented the Kolmogorov-Smirnov two sample test (Frodesen et al., 1979) available on the **stats** module of the **Scipy** package from Python (Virtanen et al., 2020). The aggregated data for all quantities considered is summarized in (Supp. Table T2).

### Analysis of ERK dynamics in long-term datasets

Long-term recordings to map ERK dynamics across the cell cycle were about 12.5 times longer and had a sampling rate reduced to about 1/5 compared to the short-term recordings (Fig. 4 Supp. 1). These qualitative differences of these data prompted for a different analysis strategy. Due to this limited time resolution, we decided to exclusively focus on the occurrence and timing of ERK pulses in the long-term datasets, and hence refer to these features as peaks.

#### Peak detection

The long-term recordings data featured both low and high frequency fluctuations. Low frequency noise created variable trends that impeded direct comparison between traces, while high frequency noise could hinder the identification of activity pulses. We used two different filtering strategies to remove fluctuations: (i) a baseline filtering that removed only low frequencies, and (ii) a band-pass filter that removed both low and high frequencies. Both methods produced similar statistics after peak detection.

In the first strategy we flattened the baseline of each trace by subtracting a low degree polynomial that follows its minima (Fig. 4 Supp. 2). To obtain this polynomial, we first identified all the local minima on each time series. We compared each value *x_i_* of the time series with its **two** neighbors to the left *x*_*i*–1_ and *x*_*i*–2_, and to the right *x*_*i*+1_ and *x*_*i*+2_. The value *x*_1_ was compared with its two right neighbors and *x*_0_ to the left, and *x*_*n*–1_ with its two left neighbors and *x_n_* to the right. We used the least squares method to fit a polynomial to the minima together with the endpoints of the trace (**numpy** (Harris et al., 2020)). Due to the variability in the traces duration and baseline, we set a trace specific polynomial degree *deg* to allow an accurate fit of the baseline while avoiding overfitting, with *deg_i_* = (2 + *m_j_*)/3, where *m_j_* is the number of minima in trace *j*.

In the second strategy, we filtered the signal by removing unwanted high and low frequencies with a band-pass filter (Fig. 4 Supp. 2). We applied a Butterworth filter with zero time and linear phase, by implementing the band-pass filter on a moving window both forward and backward in time (**scipy Signal** submodule (Virtanen et al., 2020)). We used an odd extension for the padded signal and a pad length of 15 frames, that is 3 times the number of coefficients of the Butterworth polynomials. The Butterworth filter is a band-pass square filter: it has a flat frequency response in the passband region, and rolls off towards zero in the stopband region. The order of the filter regulates the sharpness of the cutoff and we set it to 4. We chose the cutoff frequencies *f_low_* and *f_high_* in terms of the maximum frequency we can resolve with the given sampling rate. We chose low and high stopband frequencies in terms of the Nyquist frequency, *f_low_* = 0.025 *f_nyq_* and *f_high_* = 0.6 *f_nyq_*, with *f_nyq_* = 0.5 *f_s_* = (1/210) Hz for a sampling frequency *f_s_* = (1/105) Hz.

We determined the local maxima by comparison of neighboring values. We compared each value 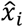 of the time series with its neighbours 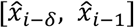 and 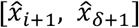, where 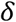 is a free parameter of the method that determines the minimum time interval between peaks that we could resolve. We reduced the range of comparison until reaching 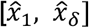 for the initial value 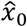, and 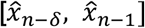 for the final value 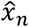. We set *δ* = 2 frames, which allowed us to resolve ERK-dependent peaks that are at least 3.5 minutes apart.

#### Threshold analysis protocol

To remove spurious low amplitude peaks, we filtered peaks with a KTR signal threshold value *I_th_*. We explored how the number of peaks changed in *Fgf4* mutant in N2B27 (negative control) and wild type cells growing in serum + LIF as we changed this threshold (Fig. 4 Supp. 2). We detected peaks in the two conditions for different 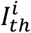 threshold values evenly spaced in the a.u. range [0, 30]. For each 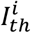, we computed the *total pulse rate*

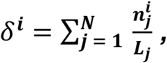

where *N* is the total number of cells of each condition, 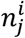 is the number of detected pulses for this threshold value 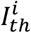 and *L_j_* is the total length of the time series of cell *j.* We normalized pulse rate to the total pulse rate *δ*^0^ at 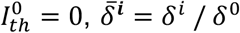 (Fig. 4 Supp. 2). This normalized pulse rate decreased with increasing the threshold values both in the negative control and the wild type. The negative control pulse rate decays much faster, reaching 0.5 while wild type values are still around 0.9. Thus, wild type genuine peaks can be distinguished from the background fluctuations in the control. We set a threshold value *I_th_* for which 1% of all the local maxima were classified as peaks in the negative control, that is 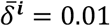. This condition results in threshold values *I_th_* = 24 for the frequency filtering strategy and *I_th_* = 25 for baseline filtering strategy (Fig. 4 Supp. 2.). We chose the baseline filtering strategy to analyze the data shown in Fig. 4, Fig. 4 Supp. 1 and Fig. 4 Supp. 3.

#### Error estimation in pulse rate maps

Being 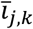 the contribution of vector 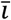 to (*T_b,j_, T_e,k_*), we interpreted each element of every 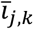 as an individual experiment with two possible outcomes 1 (success) and 0 (failure). This *T* independent experiments in (*T_b,j_, T_e,k_*) had a characteristic probability of success *p* ∈ [0,1]. Then, the probability of obtaining 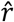 numbers of success in the *T* independent experiments in (*T_b,j_, T_e,k_*) is determined by the binomial distribution 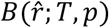.

We are interested on estimate the relative number of successes in *T* trials 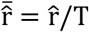. Then, the maximum likelihood estimator for 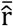 is given by 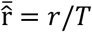 and its variance 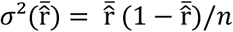 (Frodesen et al., 1979).

Understanding the *T* number of trials as a time interval, the maximum likelihood estimator of the relative number of successes is the previously defined pulse rate, that is number of peaks per unit time. Then, we estimated the pulse rate in each subspace (*T_b,j_, T_e,k_*) (Fig. 4F, Fig. 4 Supp. 3). The corresponding error was computed as the standard deviation in Fig. 4 Supp. 3. On this approach we assume stationarity conditions for each subspace (*T_b,j_, T_e,k_*) by assuming a constant *p* in each case. We neglected small variations in *p* because we wanted to study the behavior of the previously characterized short-term dynamical activity (~7 min) in long-term cell cycle time scales (~13 h).

## Supplementary Figures

**Fig. 1 Supp. 1.**
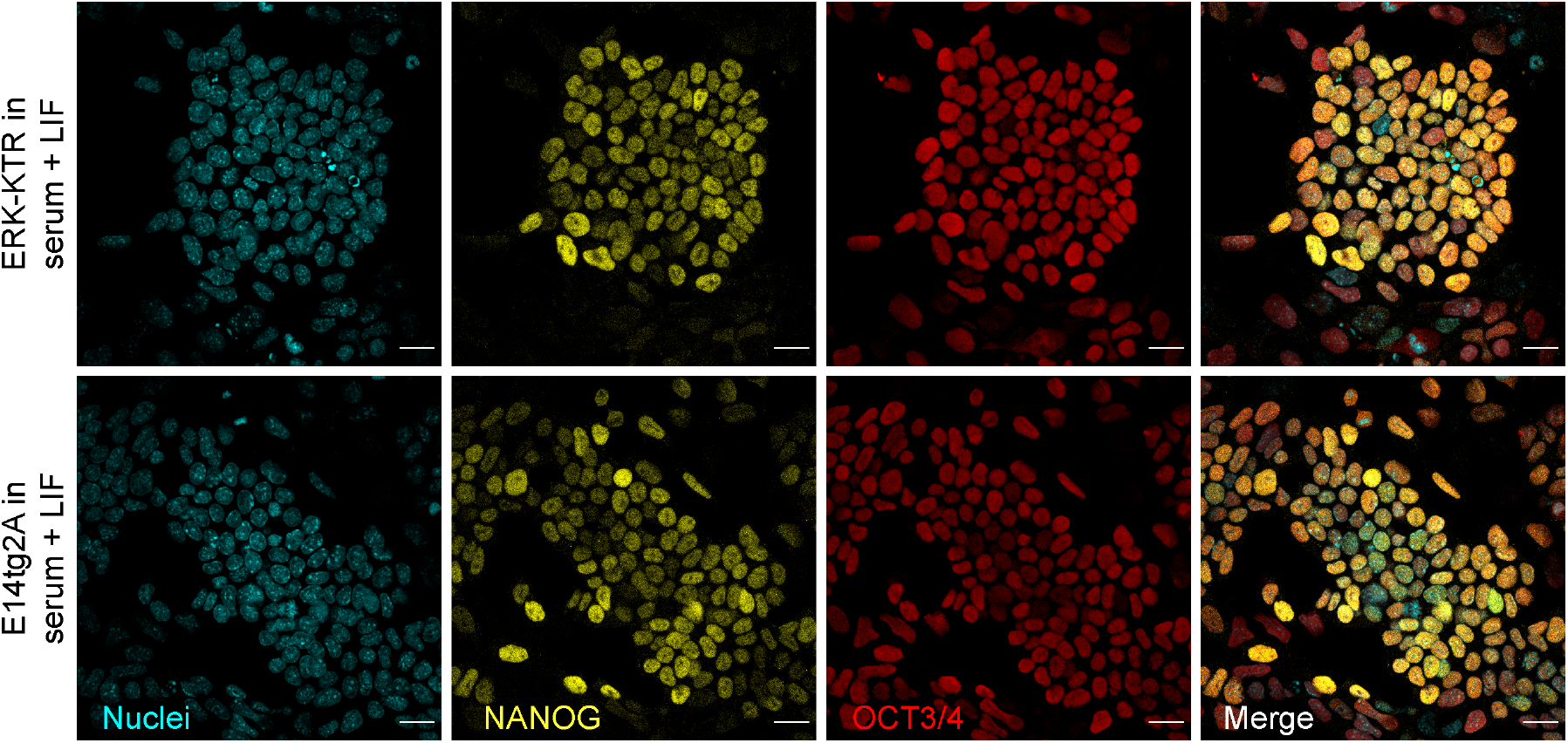
Reporter cells and parental cells express similar levels of pluripotency markers. **A.** Immunostaining of ERK-KTR mESCs (top row) and the parental E14tg2a line (bottom row) for expression of pluripotency markers NANOG (yellow) and OCT3/4 (red). Nuclei in cyan. Scale bar = 20 μm.

**Fig. 1 Supp. 2.**
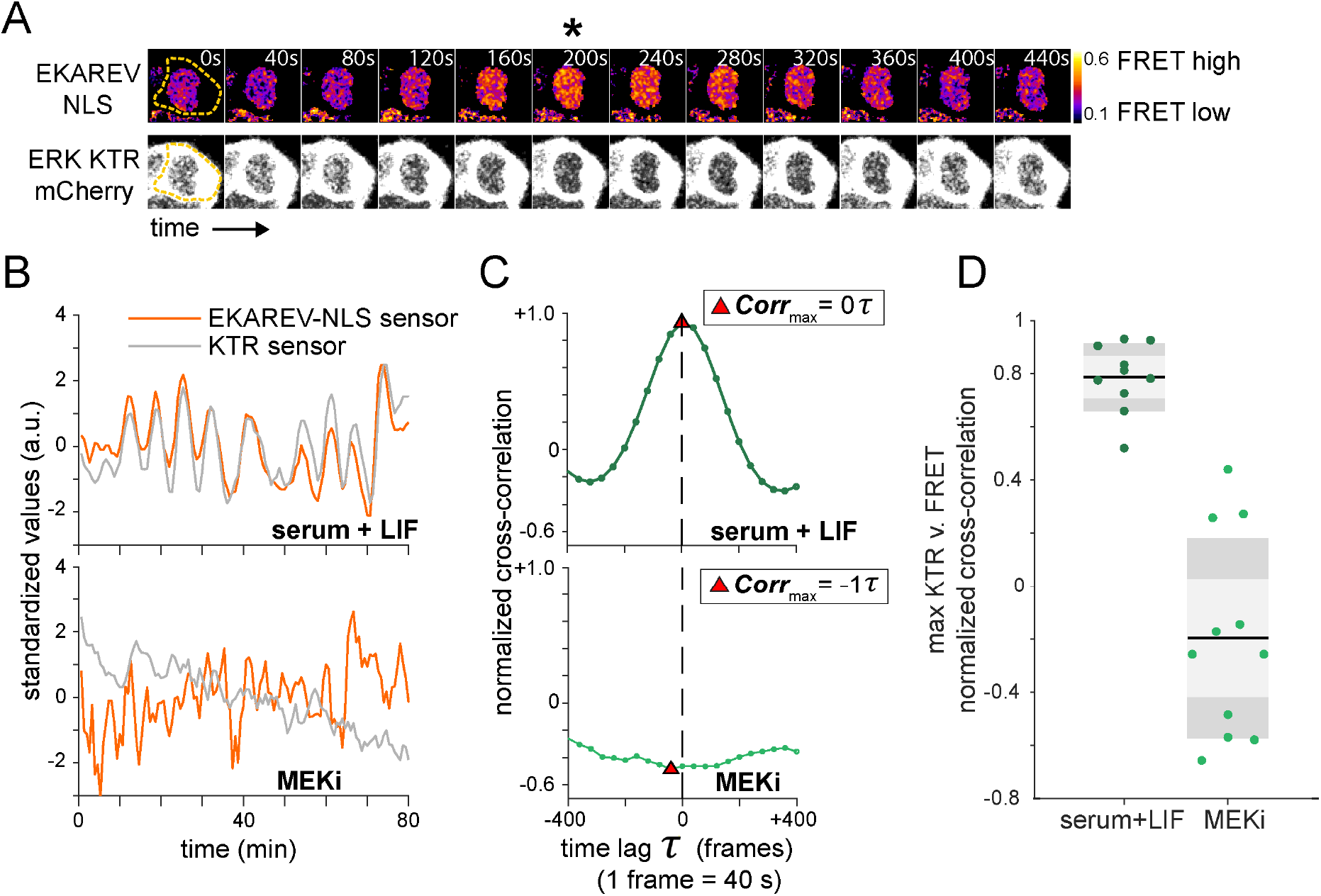
Orthogonal ERK activity sensors report similar dynamics. **A.** Stills from a movie of mESCs growing in serum + LIF medium co-transfected with both an ERK-KTR-mCherry and an EKAREV-CFP-YFP FRET reporter. Upper row shows ratiometric images of a single cell expressing the EKAREV sensor, bottom row shows images of the same cell expressing the KTR-mCherry sensor. High ERK activity detected by the FRET reporter coincides with strong nuclear exclusion of the KTR reporter (asterisk). Gamma values for the KTR montage have been adjusted to 0.86, and the image has been smoothened for the purpose of visualization only. The acquisition rate was 40 s/frame. **B.** Single cell trace of mean nuclear intensity (KTR reporter, grey) and mean FRET ratio (EKAREV reporter, orange) in the same nuclear ROI over time in the absence (top) and the presence of MEKi (bottom). FRET ratio was calculated as the ratio between donor emission and acceptor emission upon donor excitation. Traces are standardized by subtracting the mean and then dividing by the standard deviation of every individual trace. **C.** Normalized cross correlation for data shown in **B**. between traces of the different sensors as a function of time lag τ. **D.** Summary statistics of maximum cross correlation over a lag of ± 400 s between both reporters in pluripotency (N = 10 cells) and MEKi (N = 11 cells) conditions.

**Fig. 1 Supp. 3.**
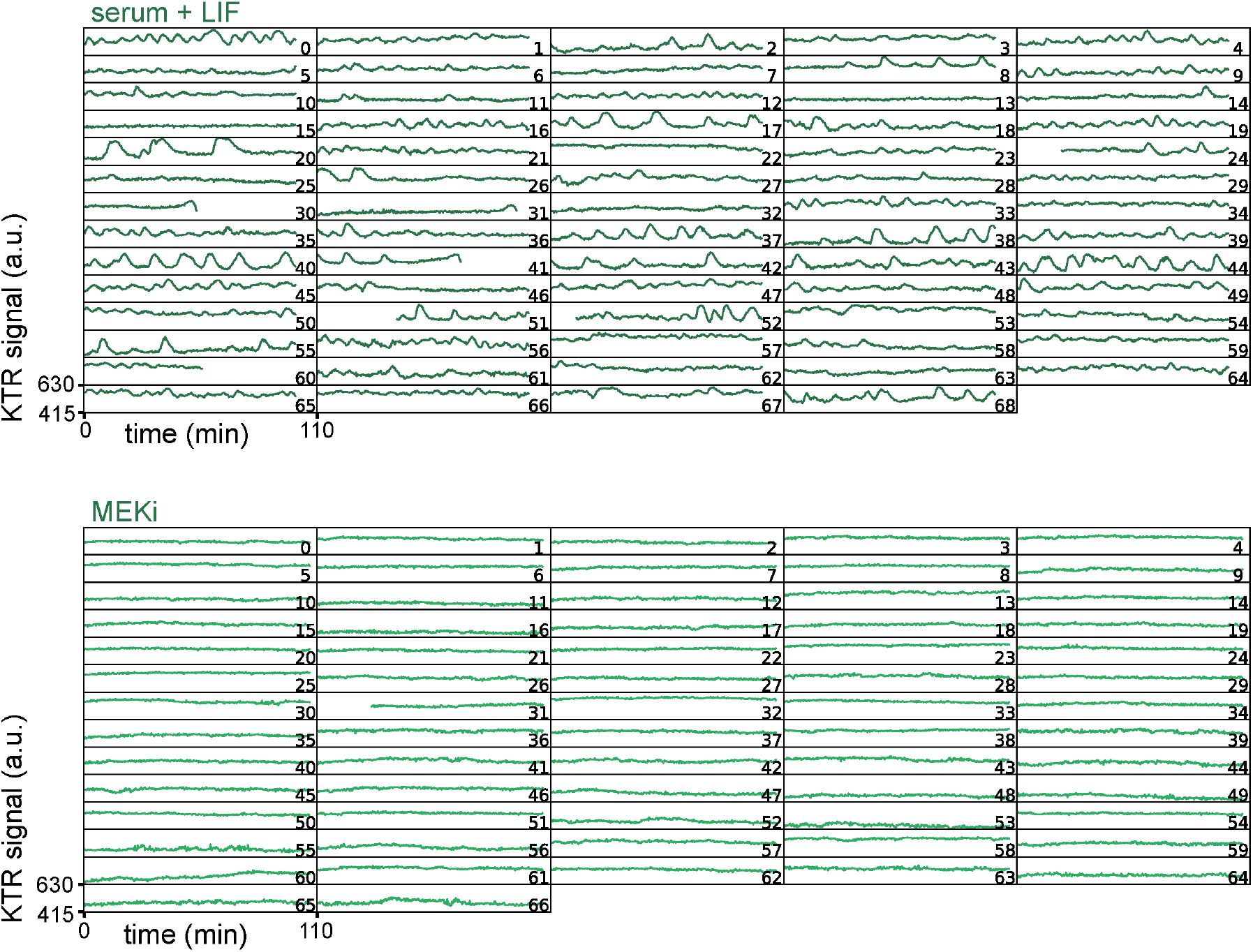
Dynamics of KTR signal reveals ERK pulsing in serum + LIF conditions. Traces of the KTR signal obtained as the mean inverted fluorescence intensity within a nuclear ROI in single cells growing in serum + LIF without (top) and with MEKi (bottom). The decrease in KTR signal at the end of the trace in cells 30, 31, 41 and 50 in the condition without MEKi is due to nuclear envelope breakdown as cells enter mitosis. This part of the trace, together with the immediately preceding peak, was trimmed for the downstream analysis. The acquisition rate was 20 s/frame.

**Fig. 2 Supp. 1.**
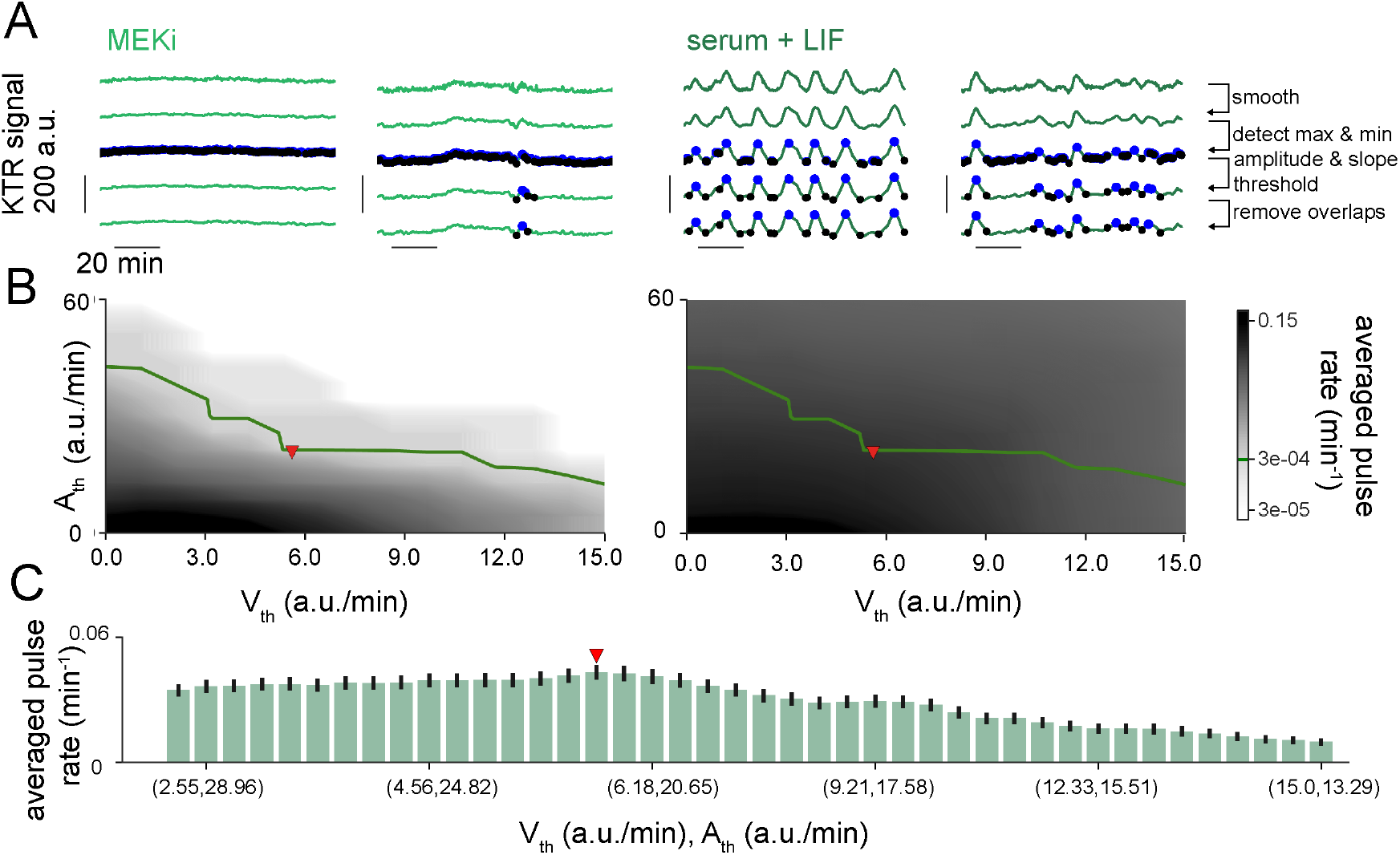
Pulse recognition and threshold analysis in high resolution time-series. **A.** Representative traces of ERK dynamical activity in single ESCs growing in serum + LIF conditions in the presence (two columns on the left) or absence (two columns on the right) of MEKi. Rows illustrate steps in the pulse recognition algorithm: First row shows raw data, second row shows smoothened traces. Blue and black dots in the third row are local maxima and minima. Fourth row shows local maxima and minima that pass the amplitude and slope thresholds. Fifth row shows identified pulses after removing overlaps. Pulses are defined by maxima and their adjacent minima. **B.** Average pulse rate as a function of amplitude and slope thresholds for cells growing in serum + LIF with (left) or without (right) MEKi. The level curve where the average pulse rate in MEKi-treated cells is 3 × 10^-4^min^-1^ (green line) was used to explore combinations of amplitude and slope threshold values in the condition without inhibitor. **C.** Average pulse rate for combinations of amplitude and slope thresholds along the red curve in cells growing in serum + LIF only. Error bar indicates SEM. Red triangle in **B**, **C** indicates parameter values used for subsequent analysis (Methods, Supp. Table T1).

**Fig. 2 Supp. 2.**
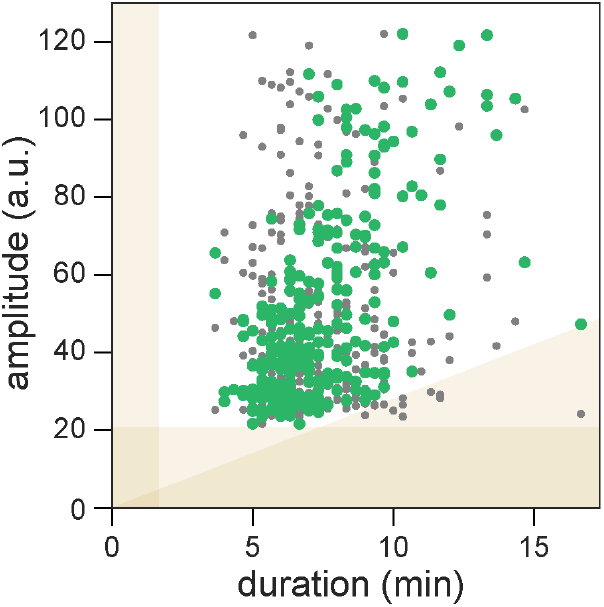
Correlation of pulse amplitude and duration in cells growing in serum + LIF. Amplitude vs. pulse duration for individual pulses (green dots). Grey dots show randomly shuffled values for comparison. Shaded yellow regions indicate the pulse recognition limits determined by the slope (yellow triangle) and amplitude (horizontal bar) thresholds in the pulse recognition algorithm, as well as the sampling resolution (vertical bar).

**Fig. 3 Supp. 1.**
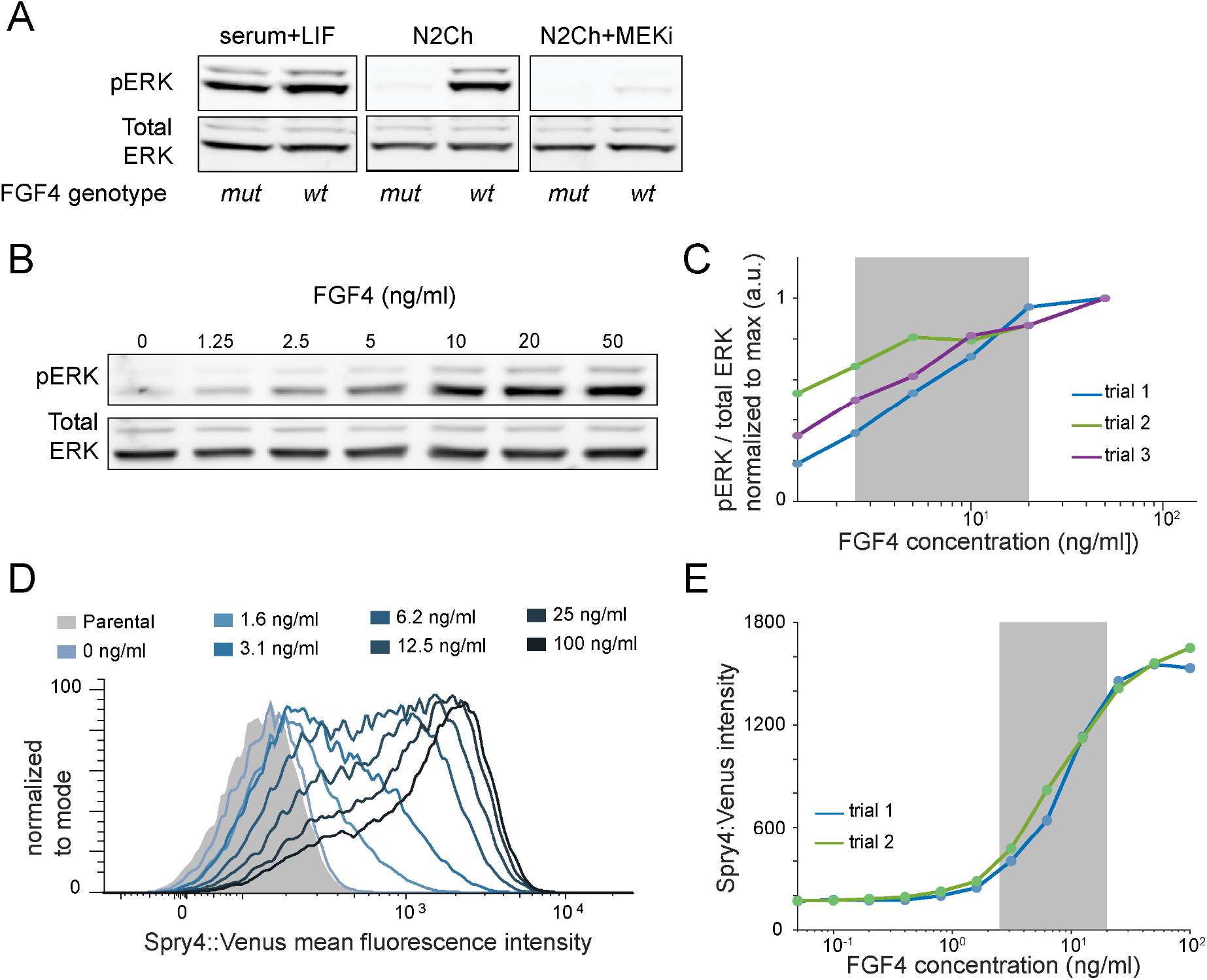
Dynamic range of signaling and transcriptional response to FGF4 dose in ESCs. **A.** Western blot for pERK and total ERK in wild type and Fgf4 mutant cells growing in the indicated media conditions. **B.** Representative western blot for pERK and total ERK in Fgf4 mutant cells treated with a range of FGF4 concentrations, with the same experimental protocol as in Fig. 3A. **C.** Quantification of western blot data from N = 3 independent experiments. **D.** Flow cytometry of Fgf4^mutant^, Spry4^H2B-Venus/H2B-Venus^ cells stimulated with a range of FGF4 concentrations as described in the methods. A non-reporter line was used as the negative control (shaded in grey). **E.** Quantification of the mean H2B-Venus fluorescence intensity from **D**. Gray box in **C** and **E** indicates the concentration range used in this study from 2.5 to 20 ng/ml.

**Fig. 3 Supp. 2.**
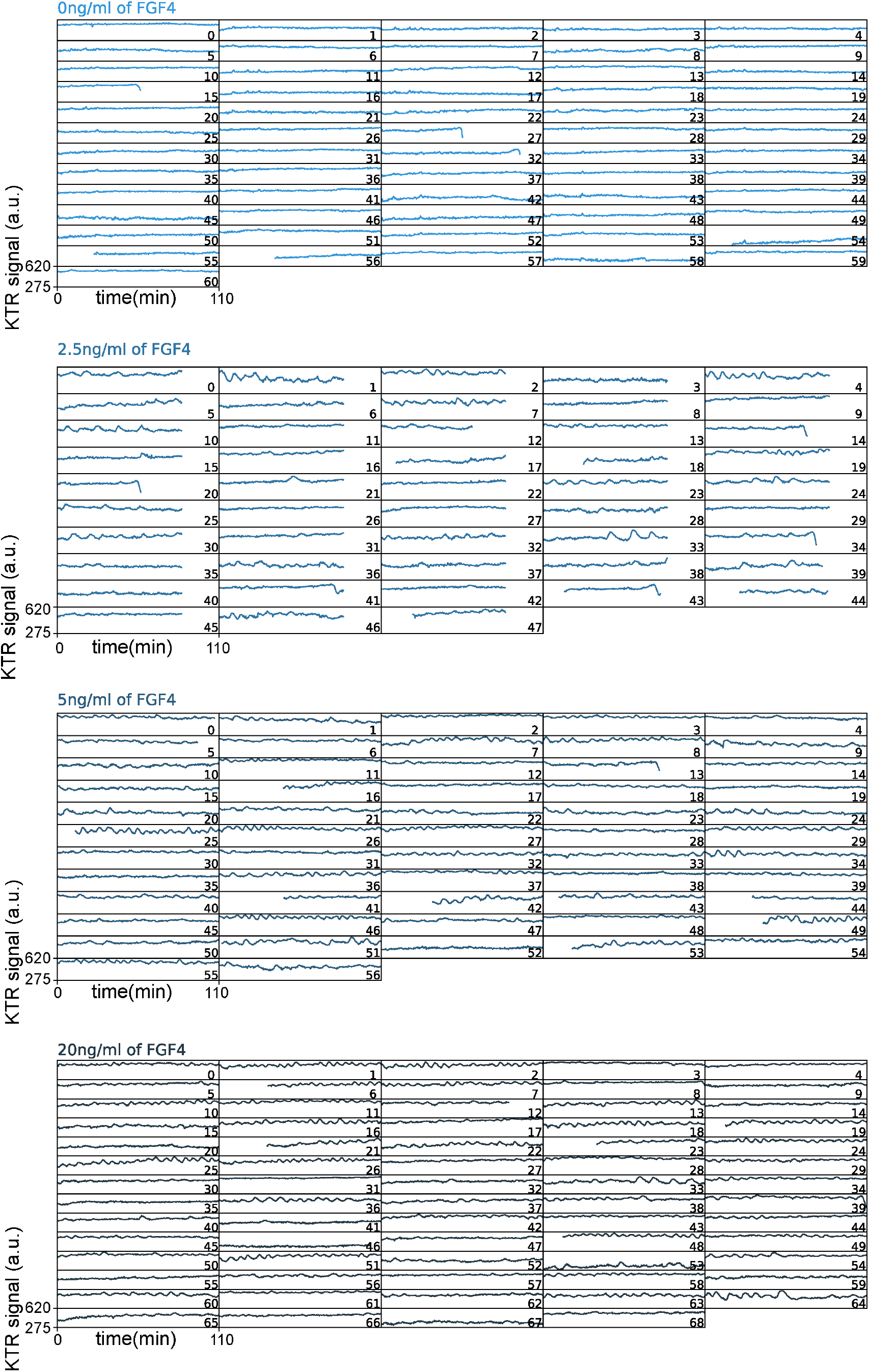
Dynamics of KTR signal at different FGF4 doses. Traces of the KTR signal obtained as the mean inverted fluorescence intensity within a nuclear ROI in single Fgf4 mutant cells stimulated with indicated doses of FGF4. The decrease in KTR signal at the end of the trace in cells 15, 27 and 32 (0 ng/ml), cells 14, 20, 34, 38, 41, 43, and 44 (2.5 ng/ml), cell 13 (5 ng/ml) and cells 8 and 39 (20 ng/ml) is due to nuclear envelope breakdown as cells enter mitosis. This part of the trace, together with the immediately preceding peak, was trimmed for the downstream analysis. The acquisition rate was 20 s/frame.

**Fig. 3 Supp. 3.**
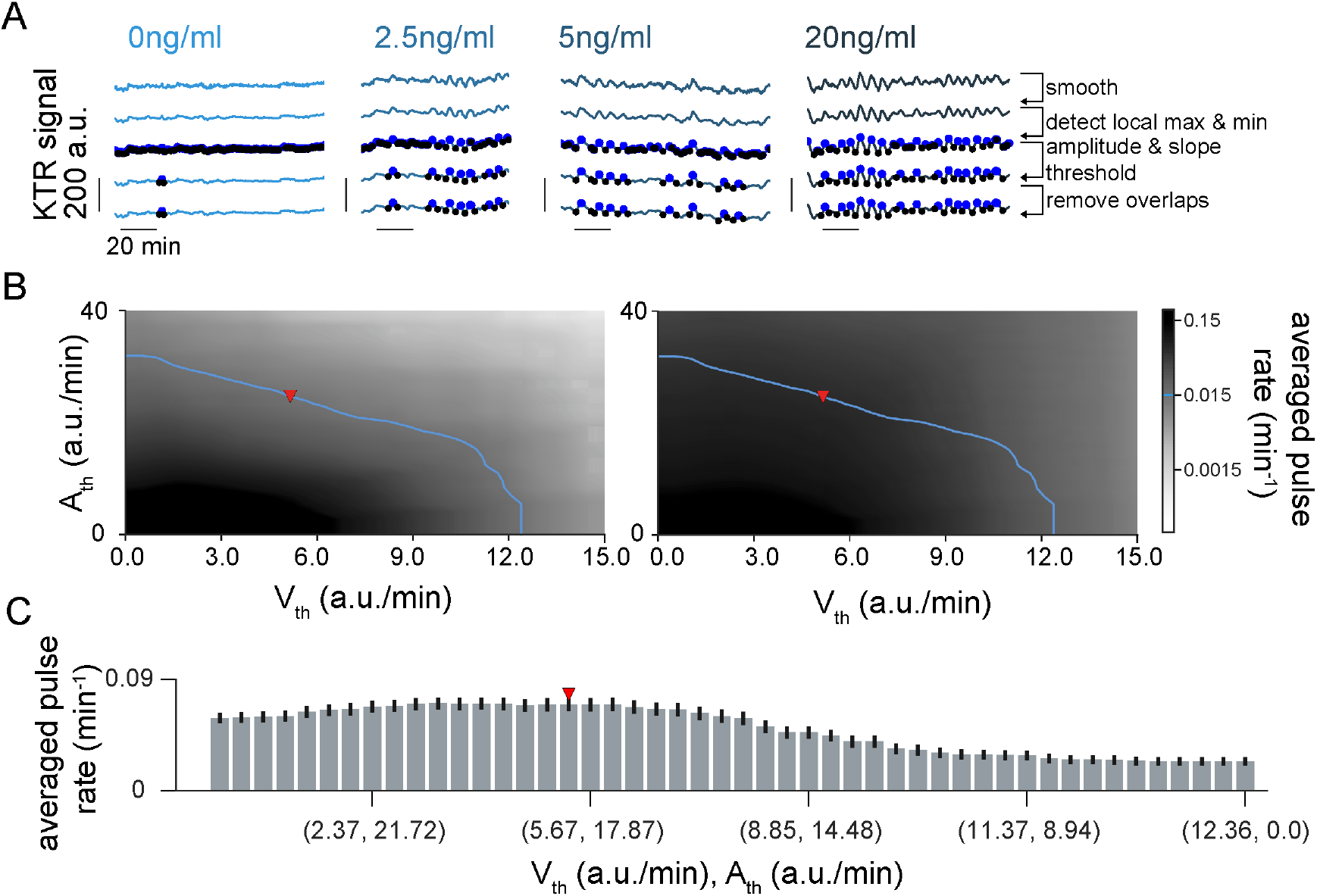
Pulse recognition and threshold analysis in FGF4 stimulation experiment. **A.** Representative traces of ERK dynamical activity in single Fgf4 mutant cells stimulated with different doses of FGF4 with colors as in Fig. 3 (columns). Rows illustrate steps in the pulse recognition algorithm: First row shows raw data, second row shows smoothened traces. Blue and black dots in the third row are local maxima and minima. Fourth row shows local maxima and minima that pass the amplitude and slope thresholds. Fifth row shows identified pulses after removing overlaps. Pulses are defined by maxima and their adjacent minima. **B.** Average pulse rate as a function of amplitude and slope thresholds for Fgf4 mutant without stimulation (left) and stimulated with 20 ng/ml FGF4 (right). The level curve where the average pulse rate in unstimulated cells is 0.015 min^-1^ (blue line) was used to explore combinations of amplitude and slope threshold values in the stimulated conditions. **C.** Average pulse rate for combinations of amplitude and slope thresholds along the blue curve in Fgf4 mutant cells stimulated with 20 ng/ml of FGF4. Error bar indicates SEM. Red triangle in **B**, **C** indicates parameter values used for subsequent analysis (Methods, Supp. Table T1).

**Fig. 3 Supp. 4.**
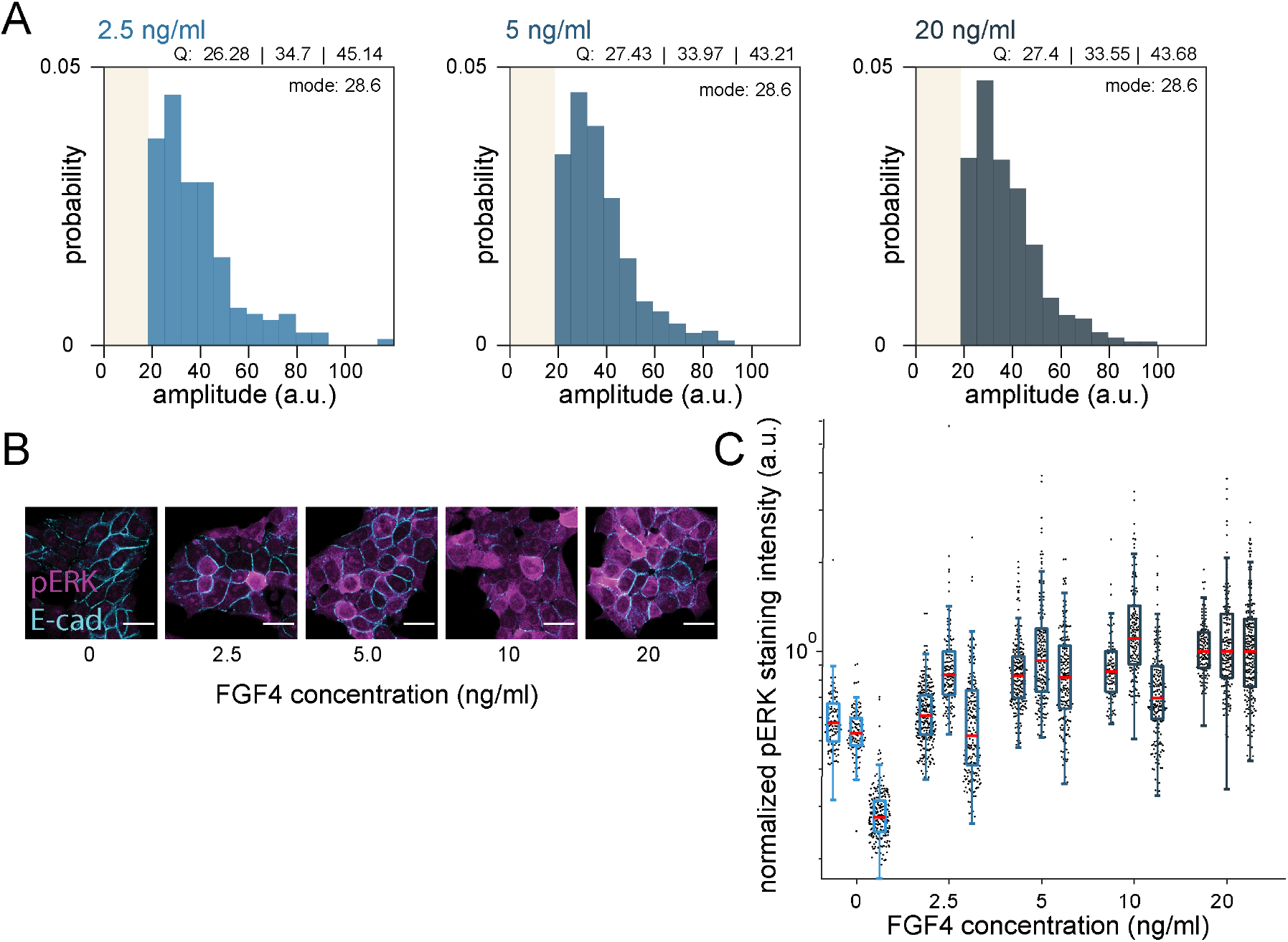
Distribution of pulse amplitudes and single cell pERK levels at different FGF4 doses. **A**. Distribution of sensor pulse amplitudes in Fgf4 mutant cells stimulated with different doses of FGF4. The number of pulses was n = 164 (2.5 ng/ml), n = 426 (5 ng/ml) and n = 544 (20 ng/ml). Pulse recognition resolution limit (yellow bar) and quartiles (Q) 25, 50 and 75 are indicated, and histograms are normalized to 1.**B**. Immunostaining of Fgf4 mutant cells for pERK (magenta) and E-Cadherin (cyan) to outline cell boundaries. Cells were treated with indicated concentrations of FGF4, with the experimental protocol depicted in Fig. 3A. Scale bar = 20 μm. **C**. Boxplot of pERK intensity in single cells stained as in **B**. Black dots represent individual cells, red bars are the median, box bounds are the 25 and 75 percentiles of the distributions, and whiskers are the 5 and 95 percentiles. Data for 3 replicates are shown for each condition. Intensity values are normalized to the median of the 20 ng/ml condition for each experiment to facilitate comparison.

**Fig. 4 Supp. 1.**
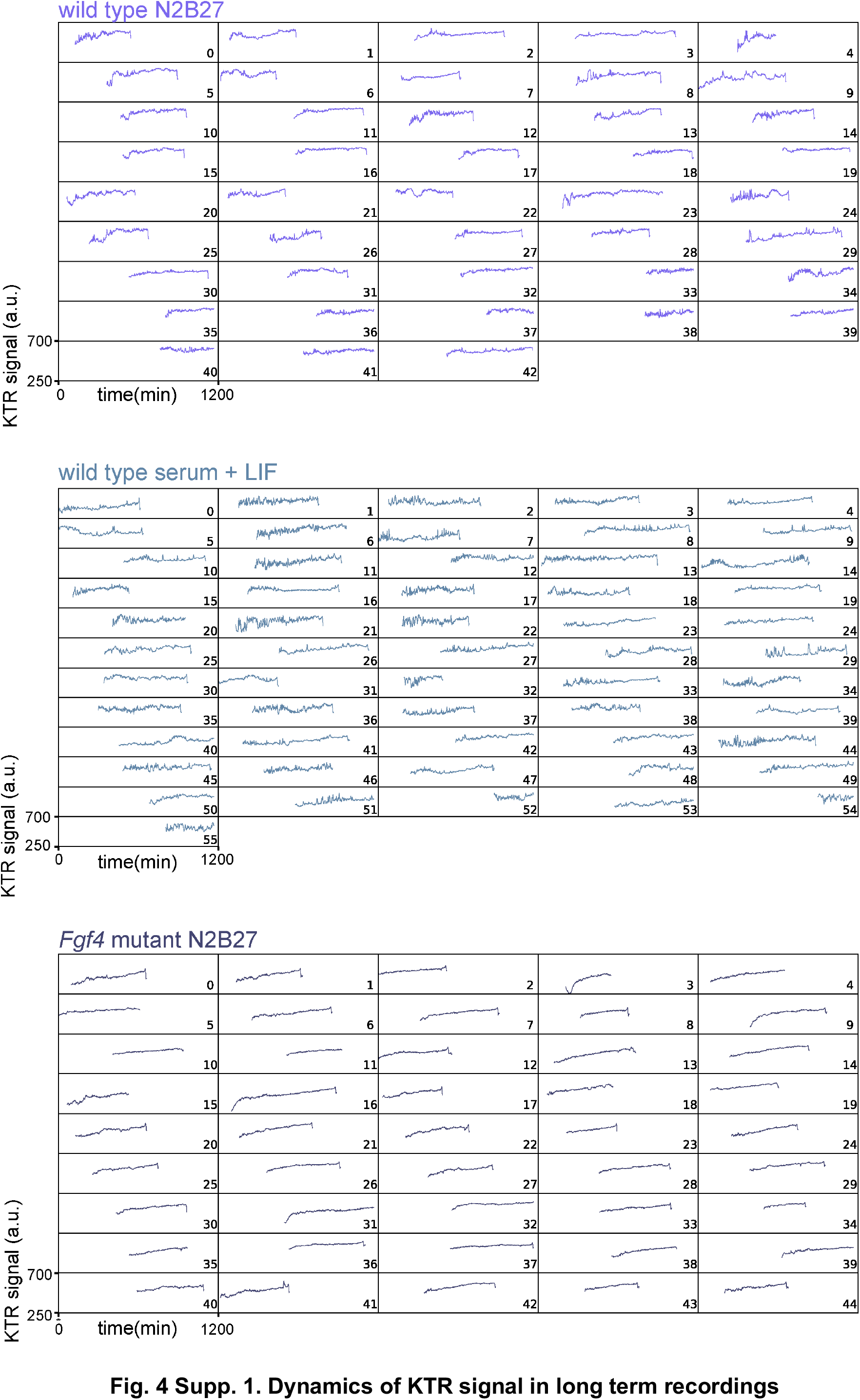
Dynamics of KTR signal in long term recordings. Traces of the KTR signal obtained as the mean inverted fluorescence intensity within a nuclear ROI in wild type cells growing in N2B27 (top), serum + LIF (middle), and in Fgf4 mutant cells growing in N2B27 (bottom). The acquisition rate was 105 s/frame. The scale of the horizontal axis represents absolute experimental time. Single cell tracks begin immediately after a cell division event and are plotted relative to absolute experimental time. Most traces end with exclusion of the sensor from the nucleus before cell division. This part of the traces, together with the immediately preceding peak, was trimmed for the downstream analysis.

**Fig. 4 Supp. 2.**
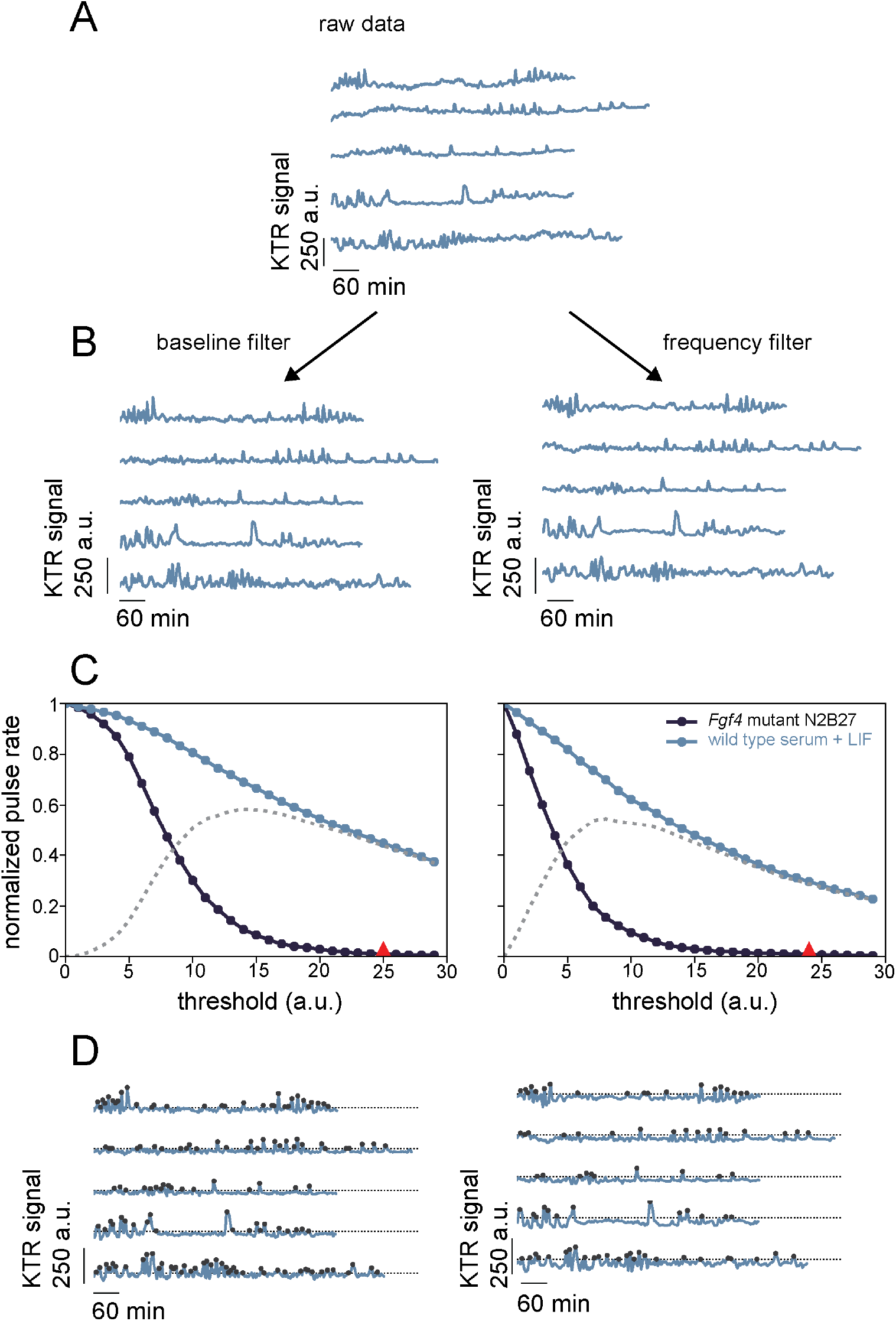
Peak detection and threshold analysis in long term time series. **A.** Representative traces of KTR signal from long term recordings in single wild type cells growing in serum + LIF. Traces have been aligned relative to the time of cell birth for this illustration. **B - D** illustrate the two filtering strategies, left column corresponds to baseline filtering and right column to band-pass filtering (Methods). **B.** Same traces as in **A** following filtering. **C.** Plot of normalized pulse rate vs. filtered KTR signal threshold to explore how the number of detected pulses depends on threshold value. Fgf4 mutant cells growing in N2B27 in dark blue, wild type cells growing in serum + LIF in light blue. The gray dotted line represents the difference of the normalized pulse rates between the experimental conditions considered. The position of the selected intensity threshold value I_th_ is marked with a red triangle. **D.** Same traces as in **B** with identified peaks (black dots). The dotted grey line in indicates the selected threshold parameter I_th_.

**Fig. 4. Supp. 3.**
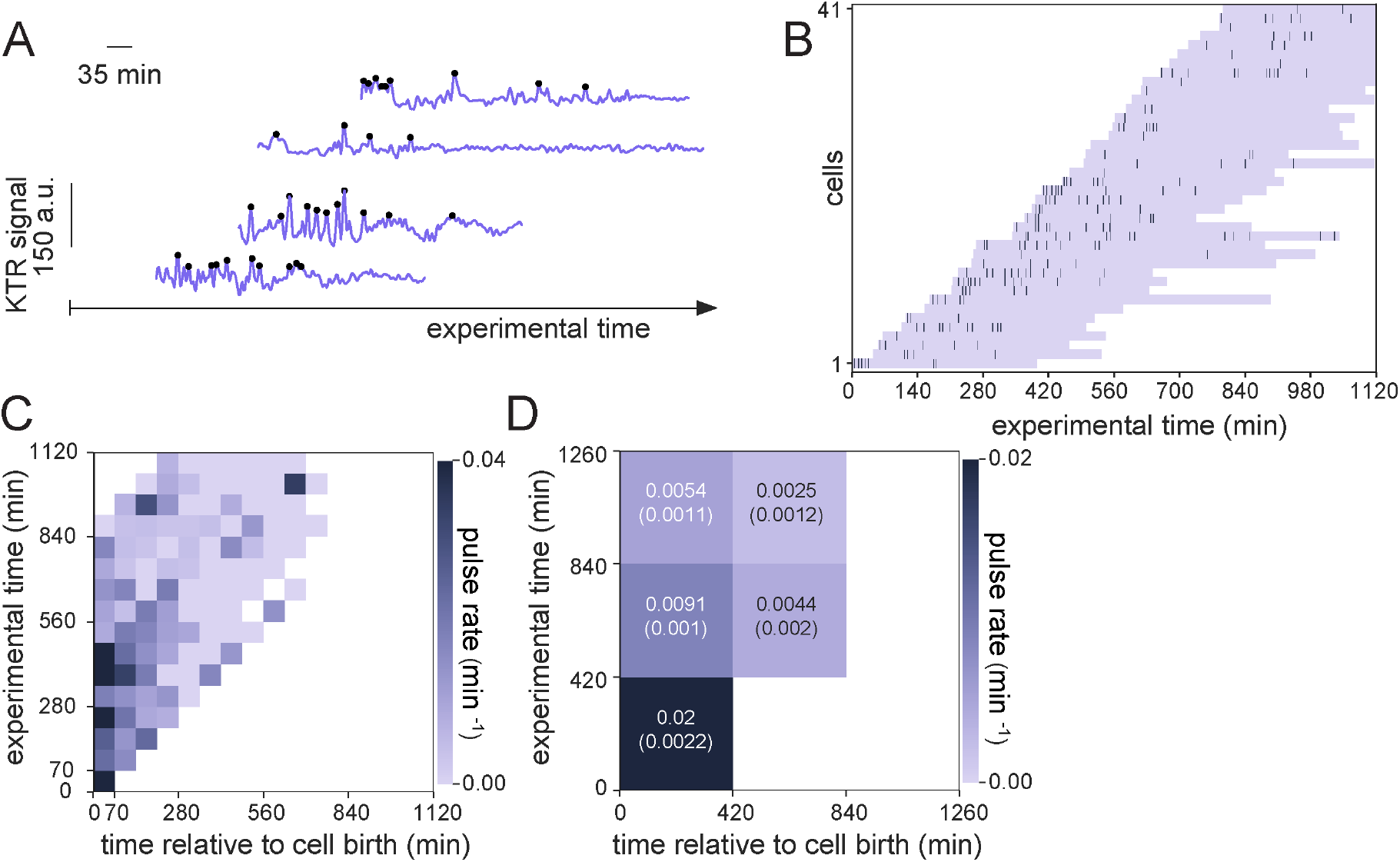
The alternative frequency filtering strategy confirms prevalent ERK pulsing early in the cell cycle. **A.** Representative traces of ERK dynamical activity from the same experiment reported in Fig. 4, now following the alternative frequency filtering strategy. Identified peaks indicated as black dots. **B.** Raster plot displaying the timing of ERK activity peaks across the cell cycle in frequency-filtered data. Rows correspond to single cells and dark bars represent peaks. Single cell tracks begin immediately after a cell division event and are plotted relative to absolute experimental time. **C.** Pulse rate map for the data shown in B. Time is discretized into 70 min bins. **D.** Coarse grained pulse rate map showing average pulse rate and its estimated error with 420 min binning.

**Fig. 4. Supp. 4.**
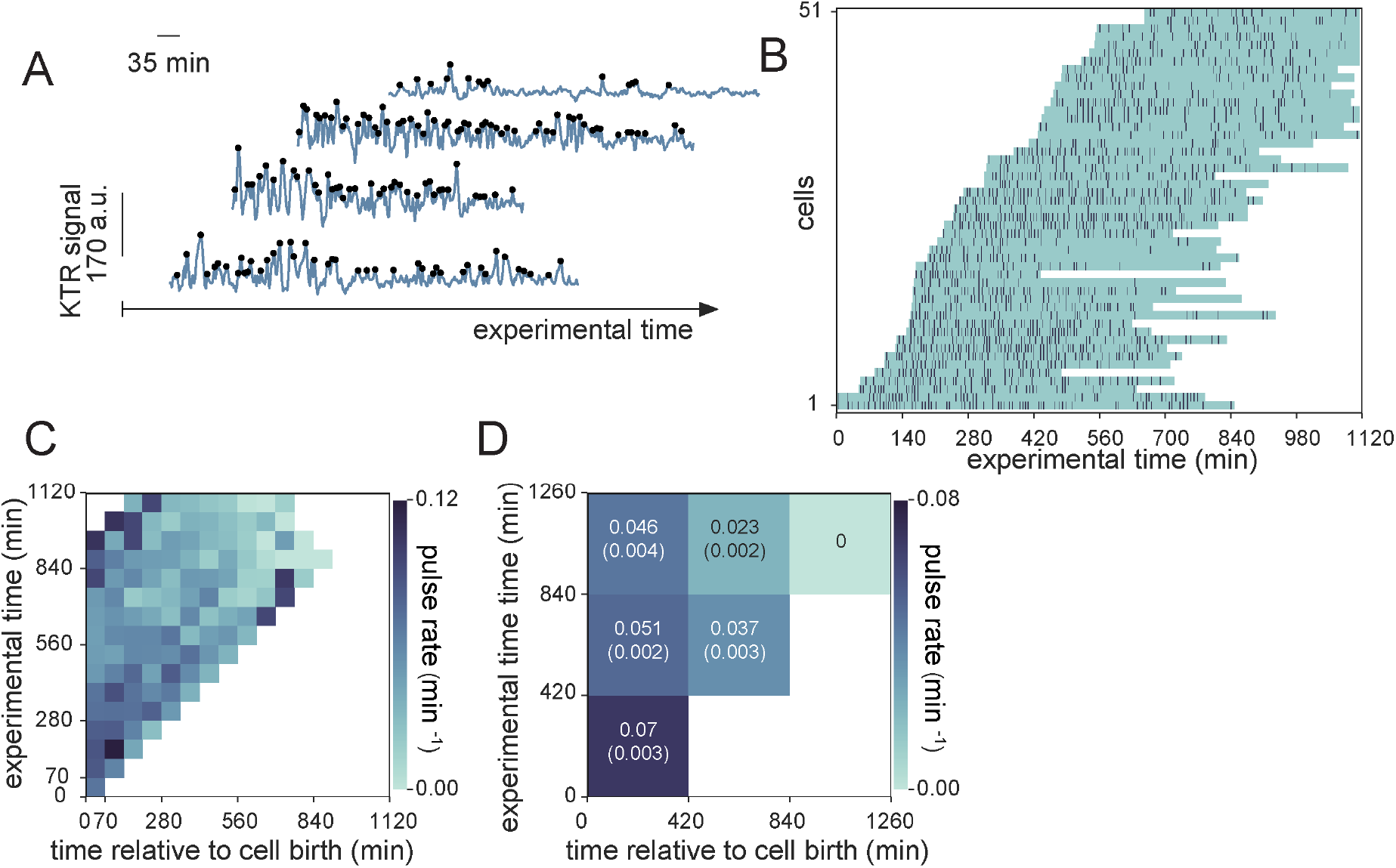
Prevalent ERK pulsing early in the cell cycle in cells growing in serum + LIF. **A.** Representative traces of ERK dynamical activity with identified peaks (black dots) in wild type cells growing in serum + LIF. Experimental protocol and baseline filtering strategy are the same as in Fig. 4. **B.** Raster plot displaying the timing of ERK activity peaks across the cell cycle in cells growing in serum + LIF. Rows correspond to single cells and dark bars represent peaks. Single cell tracks begin immediately after a cell division event and are plotted relative to absolute experimental time. **C.** Pulse rate map for the data shown in **B**. Time is discretized into 70 min bins. **D.** Coarse grained pulse rate map showing average pulse rate and its estimated error with 420 min binning.

**Supp. Table T1.** Pulse detection parameters for the threshold analysis protocol, including the amplitude and slope thresholds resulting from this protocol in the two experiments analyzed.

| Experiment | Negative control condition | Positive control condition | Exploratory parameter space | Level curve ($\delta_p^*$) | Selected amplitude threshold | Selected slope threshold |
| --- | --- | --- | --- | --- | --- | --- |
| Wild type constant stimulation | MEKi | serum + LIF | $v_{th} : (0-15) \frac{a.u}{min}$ with a $0.25 \frac{a.u}{min}$ resolution.<br>$A_{th} : (0-60)$ a.u. with a 1 a.u. resolution. | $3 \times 10^{-4} min^{-1}$ | 20.68 a.u. | $5.61 \frac{a.u}{min}$ |
| *Fgf4* mutant, different FGF4 stimulation | *Fgf4* mutant 0 ng/ml of FGF4 | *Fgf4* mutant 20 ng/ml of FGF4 | $v_{th} : (0-15) \frac{a.u}{min}$ with a $0.5 \frac{a.u}{min}$ resolution.<br>$A_{th} : (0-40)$ a.u. with a 1 a.u. resolution. | $0.015 min^{-1}$ | 18.47 a.u. | $5.16 \frac{a.u}{min}$ |

**Supp. Table T2.**
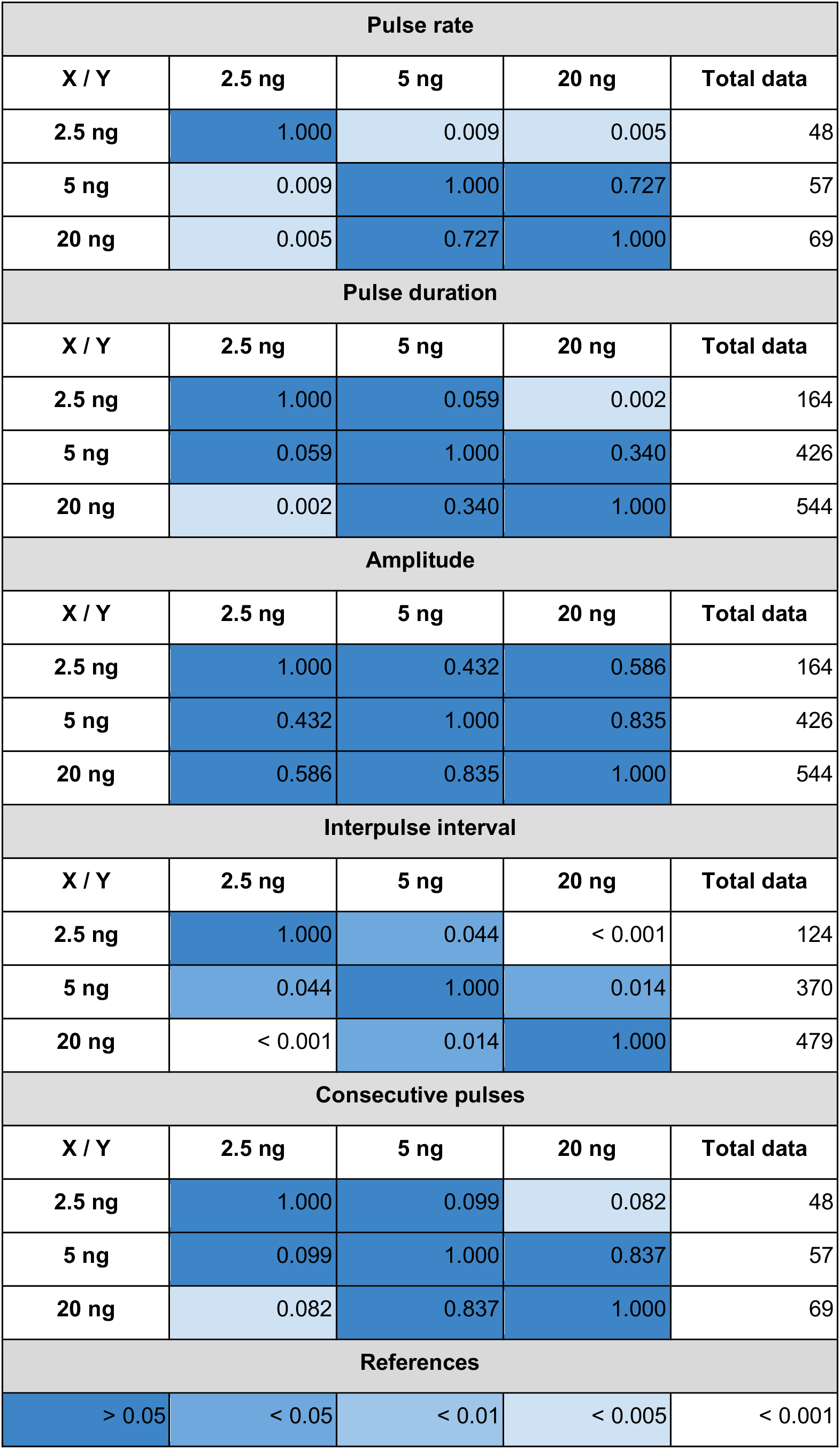
Kolmogorov-Smirnov two sample test p-value, K[x,y]. Cells values are rounded to three decimals after zero and color coded according to different p-value thresholds, the color code is given at the table bottom. The total number of data points for each condition is indicated on the rightmost column of the table. The low number of pulses at 0 ng/ml precluded statistical analysis of this condition.

